# Medial prefrontal cortex encodes but is not required to generate goal-directed actions under threat

**DOI:** 10.1101/2025.11.25.690391

**Authors:** Muhammad S. Sajid, Ji Zhou, Manuel A. Castro-Alamancos

**Affiliations:** Department of Neuroscience University of Connecticut School of Medicine Farmington CT 06001 USA

## Abstract

Adaptive behavior under threat requires deciding when to act and when to withhold action to avoid harm, often under conditions where movement, arousal, and task demand covary. Medial prefrontal cortex (mPFC) activity is widely associated with such control, yet it remains unclear whether this activity reflects causal action generation or broader evaluative processes shaped by behavioral state. Here, we combined fiber photometry, single-cell calcium imaging, mixed-effects modeling, and optogenetic inhibition to examine how GABAergic neurons in mouse mPFC represent cues, actions, and outcomes during a series of learned avoidance tasks of increasing complexity that promote cautious responding. By explicitly controlling for baseline activity and movement, we show that much apparent task-related activity in mPFC reflects movement and cue-evoked signals that are also present in a control cortical region, the visual cortex. mPFC GABAergic neurons showed little encoding of simple avoidance contingencies but broadly encoded punished outcomes. A small subset of neurons with strong movement sensitivity encoded more demanding avoidance contingencies requiring selection between action generation and deferment. For equivalent avoidance actions, distinct neuronal populations preferentially encoded either cue onset or the action. Despite this encoding, optogenetic inhibition of mPFC had minimal effects on the learning or performance of the different contingencies. These findings reveal a dissociation between neural encoding and causal necessity, indicating that mPFC GABAergic activity primarily reflects evaluative and contextual aspects of cautious avoidance behavior rather than direct control of action execution.

**Significance statement:** Avoiding danger often requires deciding when to act and when to hold back. The dorsomedial prefrontal cortex (mPFC) is widely assumed to support this control, yet the contributions of its neurons have remained unclear. Using neural population and single-neuron recordings together with targeted inhibition in mice performing learned threat-guided tasks, we show that GABAergic mPFC neurons are activated by cues, actions, and outcomes, but are not required for executing the behavior itself. These findings suggest that the mPFC primarily evaluates and contextualizes threat-motivated actions rather than generating them, highlighting a common mismatch between neural encoding and causal necessity.

### Synopsis for Reviewers

Note the following **organizational** features:

- Given the hierarchical structure of the experiments, all analyses use linear mixed-effects models, with sessions nested within mice as random effects. Models include relevant covariates, such as movement and baseline activity, to disentangle their contributions from task contingency-related effects.
- To facilitate reading flow, statistical details supporting the Results are reported in the figure legends. Additional statistical values can be provided or relocated to the main text if preferred.
- Figures and their legends are placed adjacent to the corresponding results to improve readability, but high-resolution versions are also at the end. In all population plots, the symbols, and traces are Mean±SEM. If error bars are not visible, they are smaller than the symbol and trace. We can adjust this as requested.
- Several analyses are presented as supplemental figures to streamline the main narrative; figure order and presentation can be adjusted as requested.

The **Results** are organized into six segments:

1. **mPFC and visual cortex GABAergic neurons are sensitive to movement.** Using fiber photometry, we show that GABAergic neurons in both mPFC and a control cortical region, visual cortex (VI), exhibit strong sensitivity to movement (Fig. 1; Fig. 1–S1). These findings motivated the inclusion of movement and baseline activity as covariates in neural analyses to control their effects.
2. **Behavioral performance across a series of avoidance tasks.** Mice were trained in tasks of increasing difficulty (outlined in Fig. 2A). Animals were first exposed to three neutral cues (noUS), followed by cues predicting different contingencies. In AA19, CS1 signals active avoidance, and then in AA39, CS2 signals passive avoidance. Behavioral performance is shown in Fig. 2, associated movement traces in Fig. 2–S1, and mixed-effects models of movement dynamics in Fig. 2–S2.
3. **Fiber photometry reveals limited encoding of simple avoidance but encoding of aversive outcomes.** Fiber photometry recordings from mPFC and VI during the tasks show that mPFC GABAergic neurons do not robustly encode simple avoidance contingencies but exhibit sensitivity to more complex task demands and punished errors. Analyses also tested neural responses to unsignaled aversive stimulation. Results are shown in Fig. 3, with photometry traces in Fig. 3–S1 and a full model spanning all task phases in Fig. 3–S2.
4. **Single-cell recordings reveal movement-sensitive classes with distinct contingency encoding.** Miniscope imaging was performed in mPFC during AA19 and AA39. Neurons were classified based on their correlation with movement and then tested for contingency encoding. This revealed class-specific differences, including a subset of neurons that encode avoidance-related variables. Results for AA19 are shown in Fig. 4, and for AA39 in Fig. 4–S1.
5. **Avoidance actions segregate into distinct movement modes with dissociable neural encoding.** Avoidance actions were classified based on their movement profiles, revealing three response modes that differ in response latency and vigor, reflecting varying degrees of behavioral caution. Clustering neural activity within each mode identified neurons that selectively encode cue onset versus action execution (Fig. 5 and Fig. 5–S1).
6. **Optogenetic inhibition of mPFC has minimal effects on avoidance behavior.** Using two complementary optogenetic approaches (eArch3.0 and Vgat-ChR2), we tested the causal contribution of mPFC neurons to avoidance behavior. Inhibition produced minimal effects on learning or performance across tasks. Behavioral effects and optogenetic validation are shown in Fig. 6.

## Introduction

Adaptive behavior often requires deciding when to act and when to withhold action in response to cues that predict harm (Thorndike, 1898; Mowrer, 1939; Bolles, 1970). Initiating an action promptly to prevent harm, yet not so prematurely that it incurs risk, defines cautious behavior, such as vacillating before crossing the street. Classic and contemporary theories describe cautious behavior as a trade-off between urgency and restraint (Smith and Ratcliff, 2004; Bogacz et al., 2010; van Maanen et al., 2011; Guitart-Masip et al., 2012; Heitz and Schall, 2012; Ratcliff and Frank, 2012; Yee et al., 2022; Zhou et al., 2022), often interpreted as increased cognitive evaluation prior to decision making (Donders, 1969; Sternberg, 1969; Posner, 2005). However, most studies of cautious responding have focused on appetitive contingencies (Guitart-Masip et al., 2012; Millner et al., 2018). In contrast, experimental active and passive avoidance procedures, which require animals to act or withhold action to avoid punishment, provide a framework for studying how neural circuits support cautious behavior under threat.

The dorsomedial, including prelimbic, prefrontal cortex (mPFC) is widely thought to link predictive cues to appropriate actions while suppressing inappropriate or ill-timed ones (Ridderinkhof et al., 2004; Euston et al., 2012; Le Merre et al., 2021; Friedman and Robbins, 2022). In rodents, mPFC activity is frequently reported to correlate with cue processing, action selection, and outcome evaluation during cued tasks. GABAergic neurons are central to cortical computation and are thought to shape information flow, gain control, and behavioral flexibility within mPFC circuits (Kawaguchi and Kubota, 1997; Gupta et al., 2000). Their activity is closely coupled to excitatory neurons, and frequently modulated during decision-making and learned tasks, making them a compelling population in which to examine mPFC activity. However, the functional significance of these signals remains unresolved because there are two main challenges in interpreting this activity.

A first challenge is that neural activity during behavior is strongly shaped by movement and behavioral state. Arousal is a dominant behavioral-state category long-known to shape neural activation patterns (e.g., (Castro-Alamancos and Connors, 1996; Castro-Alamancos, 2004a, 2004b)). Across cortical and subcortical regions, a large fraction of neural variance is explained by ongoing movement, often exceeding that attributable to task variables themselves (Musall et al., 2019; Steinmetz et al., 2019; Stringer et al., 2019; Zagha et al., 2022; Hormigo et al., 2023a). Such state-related modulation is especially prominent during tasks involving cue-evoked orienting and locomotion, where movement is both a response and a confound. Without explicitly accounting for movement and baseline state, activity attributed to cue encoding or action selection may instead reflect correlated changes in behavioral dynamics.

A second challenge is that identifying neural signals that encode task variables does not establish that those signals are necessary for generating behavior. Neural activity may reflect evaluation, monitoring, or contextual processing without exerting direct causal influence on action execution (e.g., (Jonas and Kording, 2017; Krakauer et al., 2017; Shenoy and Kao, 2021; Barack et al., 2022)). This distinction between encoding and necessity is particularly important for interpreting mPFC function, where correlational evidence has often been taken to imply executive control.

To address these issues, we used two approaches. First, in mice performing a series of auditory-cued avoidance tasks of increased difficulty, we combined fiber photometry and single-cell calcium imaging with detailed movement tracking and mixed-effects modeling to quantify how mPFC GABAergic activity relates to task variables after controlling for baseline state activity and movement. To determine whether the observed signals were specific to mPFC, we performed parallel recordings in visual cortex (VI), a cortical region not expected to strongly contribute to auditory cue-guided avoidance but similarly subject to movement- and baseline-related modulation. Second, we tested causal necessity using optogenetic inhibition across dorsal, prelimbic, and infralimbic mPFC areas and VI during task performance. If mPFC activity directly generates cautious actions, we predicted that it would selectively encode action execution or deferment independent of movement and that its inactivation would disrupt learning or performance. Alternatively, if mPFC activity primarily reflects evaluative or contextual processes, we expected it to encode task contingencies without being required for task performance.

Our results support the latter view. GABAergic neurons in both mPFC and VI are strongly modulated by movement, but mPFC neurons also encode some aspects of the more challenging task contingencies after controlling for movement and baseline state. Despite this encoding, broad optogenetic inhibition of mPFC using two different methods produced minimal effects on learning or performance across tasks, comparable to inhibition of VI. Together, these findings reveal a dissociation between encoding and causal necessity, indicating that mPFC activity reflects evaluative processing during cautious behavior rather than direct control of action execution.

## Results

### mPFC GABAergic activity is sensitive to movement

To assess the population activity of GABAergic neurons in mPFC during spontaneous movement, we expressed GCaMP7f (Chen et al., 2013) via local injection of a Cre-dependent AAV in Vgat-Cre mice (Vong et al., 2011). A single optical fiber was implanted in the dorsomedial, *aka* prelimbic, area of the mPFC (Mark et al., 2018); based on corpus callosum landmarks, the prelimbic area is centered between the anterior forceps and the genu, with the infralimbic area ventral to the prelimbic area, while the orbital area lies anterior and ventral to the anterior forceps. Calcium signals were recorded using fiber photometry, as previously described (Hormigo et al., 2021, 2023b). Figure 1A (left) shows a representative trajectory of an optical fiber targeting GCaMP-expressing GABAergic neurons in mPFC. The estimated imaged volume extended ∼200 µm from the fiber tip, encompassing approximately 2.5 × 10⁷ µm³ (Pisanello et al., 2019). As a control, we performed the same procedure in the visual cortex (VI), a functionally distinct cortical area (Fig. 1A). This allowed us to establish the specificity of the changes observed in mPFC. GCaMP7f was expressed in GABAergic neurons in Vgat-Cre mice, and an optical fiber was implanted in the center of VI (Fig. 1A middle). Figure 1A (right) includes a schematic showing the lens endings for both mPFC and VI implantations based on histological reconstructions. The mPFC recordings were confirmed to be centered on the prelimbic area of mPFC, which has been previously proposed to be critically involved in avoidance behavior under threat in mice (e.g., (Jercog et al., 2021; Martin-Fernandez et al., 2023)).

**Figure 1.**
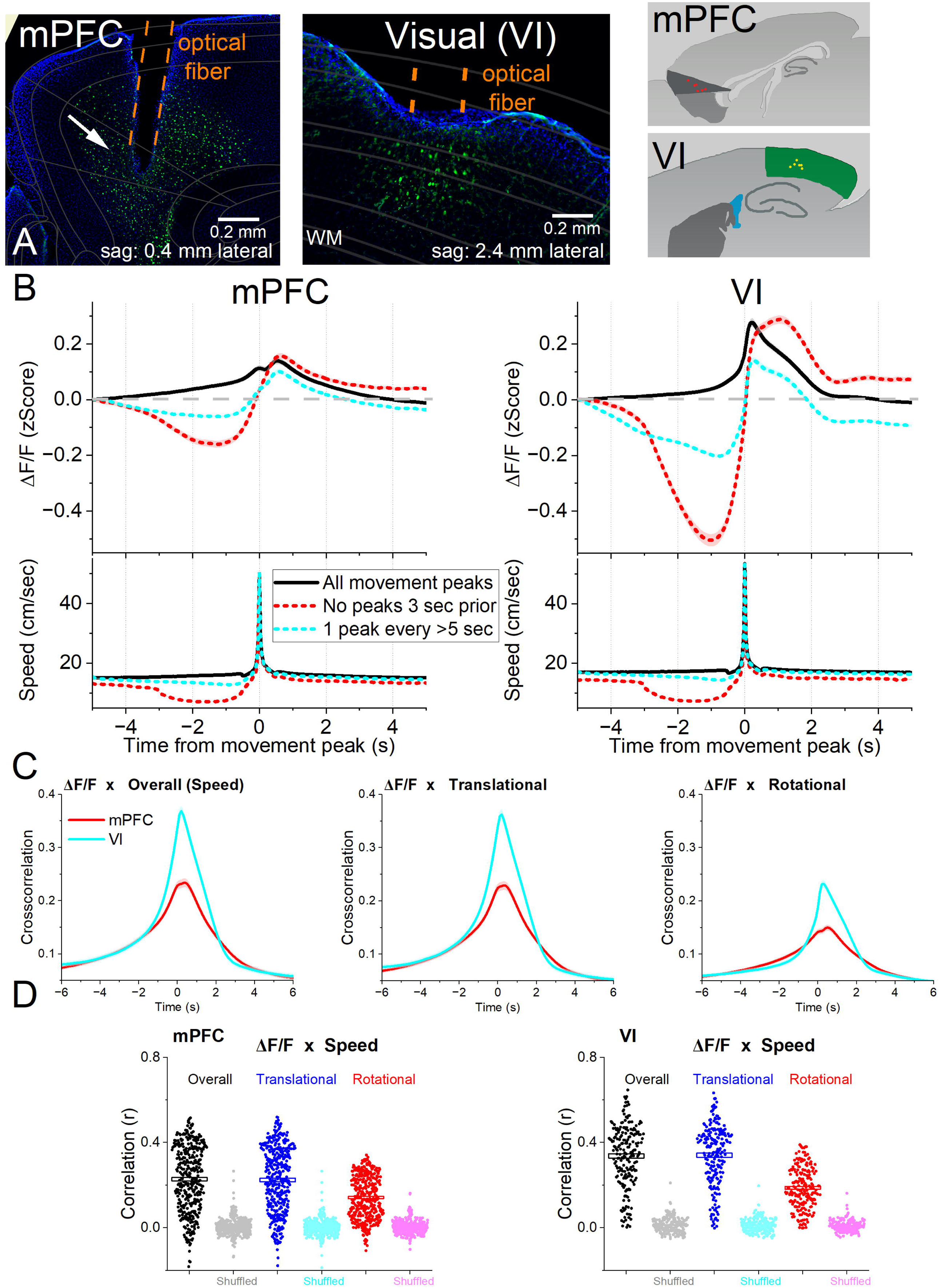
Calcium imaging fiber photometry reveals that both mPFC and VI GABAergic neurons activate during spontaneous exploratory movement. ***A***, Parasagittal sections showing the optical fiber tract reaching mPFC or VI with GCaMP7f fluorescence expressed in GABAergic neurons around the fiber ending. The sections were aligned with the Allen brain atlas CCF. The right panels show schematics of the locations of optical fiber endings (dots) in mPFC (dark gray is prelimbic area and ventral is orbital and infralimbic) and VI (green). WM, white matter. Lateral is from the midline. Left, is anterior in all panels. ***B***, ΔF/F calcium imaging time extracted around detected spontaneous movements recorded in mPFC (left) or VI (right). Time zero represents the peak speed of the movement. The upper traces show ΔF/F mean±SEM of all movement peaks (black), those that had no detected peaks 3 s prior (red), and peaks taken at intervals >5 s (cyan). Dashes are added for visualization of overlapping traces. The lower traces show the corresponding movement speed for the selected peaks. Note that the peak speeds at zero fully overlap. Linear mixed effects model contrasts: (mPFC vs VI [−3 to −0.5 s]: all peaks t(10)=0.39 p=0.70; no peaks 3 s prior: t(10)= 3.43 p=0.0064; peaks > 5 s: t(10)= 3.2 p=0.009). ***C***, Cross-correlation between movement and ΔF/F for the overall (left), translational (middle) and rotational (right) components in mPFC (red) and VI (cyan). ***D***, Per session (dots) and mean±SEM (rectangle) linear fit (correlation, r) between overall movement and ΔF/F in mPFC (left) and VI (right), including the rotational and translational components. The lighter dots show the linear fit after scrambling one of the variables (shuffled). (n=7 mice in mPFC, n=5 mice in VI) All traces and averaged symbols in the paper are mean±SEM. If SEM is not visible, it is smaller than the trace/symbol thickness.

In freely moving mice (n=7 mice in mPFC and n=5 mice in VI), we recorded continuous calcium fluorescence (ΔF/F) and spontaneous movement from three markers on the head to derive translational and rotational components, as the animals explored an open arena. To examine the relationship between movement and GABAergic neuron activation in the mPFC and VI, we identified spontaneous movement peaks and aligned the fluorescence traces to those events, following procedures established in our previous studies (Hormigo et al., 2023b; Zhou et al., 2023, 2024). Movement peaks were categorized into three groups. The first category included all detected peaks (Fig. 1B, black traces), the second category captured movement onsets from immobility, defined as peaks with no movement detected in the preceding 3 seconds (Fig. 1B, red traces), and the third category included peaks separated by at least 5 seconds (Fig. 1B, cyan traces), minimizing the impact of closely spaced movements and isolating activity related to increases from an ongoing baseline. Per session, the peaks in the second category represent ∼23% of the total peaks in the first category, while the peaks in the third category represent ∼31% of the total peaks. In all three categories, GABAergic activity in both regions increased significantly around movement compared to baseline. Although our goal was not to compare both cortical regions, VI neurons tended to exhibit stronger activation than mPFC neurons in response to equivalent movement. For instance, VI neurons showed a sharper drop in activation during the period of movement cessation preceding onset (Fig. 1B, red traces; No peaks 3 sec prior), suggesting that VI activity more closely tracks ongoing motion.

To explore these relationships further, we computed cross-correlations between movement (speed) and GABAergic activity in the mPFC and VI groups (Fig. 1C). Movement was significantly correlated with both mPFC and VI activity with the latter showing a stronger relation. When movement was decomposed into translational and rotational components, both were strongly correlated with mPFC and VI activity, with translational movement contributing most strongly to the overall correlation (Fig. 1C). We next performed a linear fit between ΔF/F and movement (speed); both ΔF/F and speed signals were segmented into consecutive 200 ms windows across the entire recording session, and the linear relationship was computed across these paired segments. This analysis revealed a strong positive linear correlation between mPFC or VI activity and both translational and rotational movement, with correlations consistently stronger in VI than in mPFC (Fig. 1D). Shuffling one variable in each pair abolished the observed correlations (Fig. 1D), confirming that the relationships do not reflect chance alignment.

We next asked whether mPFC and VI GABAergic activity during movement depends on the direction of head turns relative to the recording site. Figure 1-Suplement 1 shows head turns in the contraversive (cyan) and ipsiversive (red) directions, aligned to the corresponding ΔF/F signals recorded from mPFC (Figure 1-Suplement 1A) and VI (Figure 1-Suplement 1B) neurons across the three categories of movement events. To assess direction-dependent responses, we compared the mean amplitude, peak amplitude, and peak timing of the ΔF/F signals for equivalent ipsiversive and contraversive turns using linear mixed-effects models with Group (mPFC, VI) and Turn Direction (ipsi-, contraversive) as fixed effects factors and sessions nested within mice as random effects. In the mPFC, neuronal activity did not differ between ipsiversive and contraversive turns across any condition, indicating that GABAergic neurons in this region do not encode head turn direction (Figure 1-Suplement 1C). In contrast, VI neuron activity showed modest direction selectivity (Figure 1-Suplement 1D). For all turns, contraversive movements elicited significantly greater ΔF/F responses. This effect remained significant for isolated turns, but not for turns following a period of quiescence, suggesting that direction selectivity emerges in VI more clearly during ongoing movement.

Together, these results indicate that GABAergic neurons in both mPFC and VI are sensitive to movement. In addition, mPFC GABAergic neurons are insensitive to head movement direction, whereas VI GABAergic neurons have a modest preference for contraversive turns, particularly during sustained or ongoing movement.

### Cue Driven Action Selection Tasks

Having established that cortical GABAergic neurons in both mPFC and VI are sensitive to movement, we then evaluated their activation during cued goal-directed behaviors. Mice performed three successive tasks—noUS, AA19, and AA39—in a shuttle box (Fig. 2A) during calcium imaging fiber photometry of mPFC and VI neurons (n=7 mice in mPFC, n=5 mice in VI). In every task, mice were presented with three auditory tone CSs—CS1 (8 kHz), CS2 (4 kHz), and CS3 (12 kHz)—each delivered randomly at low, medium, and high intensity levels (∼65-87 dB). We used three tone intensities to increase the likelihood of task errors, as mice are very proficient at performing these tasks. Low-intensity cues are harder to detect, whereas high-intensity cues are harder to discriminate because loud sounds produce broader cochlear activation, reducing frequency specificity (Malmierca and Ryugo, 2012). In every task, each CS trial was followed by a random intertrial interval during which crossings (ITCs) had no consequence (Fig. 2A).

**Figure 2.**
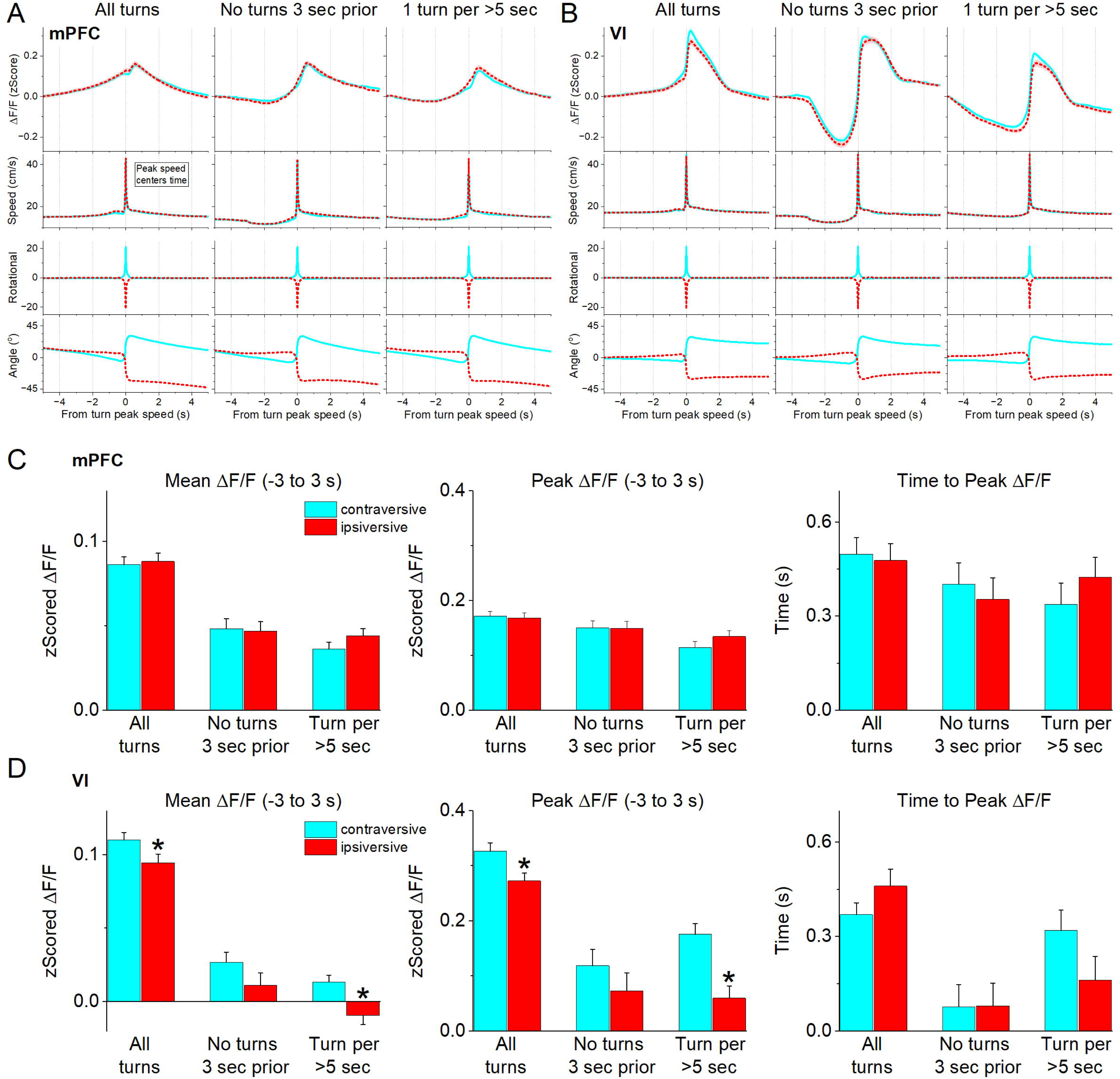
Behavioral performance in the cue-driven action tasks. ***A***, Arrangement of the shuttle box and tasks. The *top table* details the sequence of four tasks and the contingencies for each of the three CSs per task. Note that the tones do not apply to the final *unsignaled US* task (US, aversive unconditioned stimulus consisting of foot shock plus white noise). In this task, the full US, the foot shock component, or the white noise component are presented randomly and elicit a fast escape crossing (see text for further details). In the other three tasks (noUS, AA19, and AA39), three auditory tone conditioned stimuli (CS1–CS3) were presented randomly at three intensities (low, medium, high). During noUS, all CSs were neutral and crossings terminated the CS without consequence (Ig, Ignore contingency). During AA19, CS1 added the Active Avoid (AA) contingency; CS2 and CS3 remained neutral. During AA39, CS2 added the Passive Avoid (PA) contingency, while CS1 retained the AA contingency and CS3 remained neutral. The *bottom table* describes the trial structure for the three task contingencies (AA, PA, Ig). Each trial consists of a random intertrial interval (ITI), followed by a CS action interval. In the AA contingency this is followed by the US, if the mouse did not cross (actively avoid), which causes a fast crossing (escape, AA error). The PA contingency requires the mouse not to cross and applies a short US (0.5 s) if the mouse crosses (PA error). ***B***. Behavioral performance across three sequential procedures: noUS, AA19, and AA39. Panels show percentage of actions (top), action latency (middle), and intertrial crossings (ITCs; bottom). Data are shown for combined mPFC and VI groups after confirming no group effect (χ²(1) = 0.95, p = 0.32). Other model factors (Task, CS, Tone Intensity) showed significant main effects and interactions (p < 0.0001). (Models performance: Actions: R^2^_m_ = 0.44; R^2^_c_ = 0.66 | Action Latency: R^2^_m_ = 0.1; R^2^_c_ = 0.25 | ITCs/Trial: R^2^_m_ = 0.05; R^2^_c_ = 0.75) **noUS**. The number of spurious actions did not differ across CS1–CS3. Within each CS, tone intensity significantly modulated responding, with higher action rates at high compared to low intensity (t(2316) > 2.4, p < 0.05). CS2 at high intensity elicited more actions than CS2 at medium intensity (t(2315) = 5.6, p < 0.0001). **AA19**. Action rates during CS1 were higher than during noUS for all intensities (low: t(1585) = 13.02, p < 0.0001; medium: t(1575) = 15.50, p < 0.0001; high: t(1585) = 16.89, p < 0.0001). Actions during CS2 and CS3 did not differ from noUS at low or medium intensity but increased at high intensity (CS2: t(1594) = 6.97, p < 0.0001; CS3: t(1585) = 3.93, p = 0.012). Within AA19, action rates during CS1 exceeded those during CS2 (t(2315) = 24.5, p < 0.0001) and CS3 (t(2315) = 25.4, p < 0.0001), while CS2 and CS3 did not differ. Action latencies did not differ between noUS and AA19 for any CS. **AA39**. Action rates during CS1 exceeded those during CS2 and CS3 (CS1 vs. CS2: t(2316) = 21.45, p < 0.0001; CS1 vs. CS3: t(2316) = 28.6, p < 0.0001). Compared to AA19, overall responding decreased across all CSs (CS1: t(574) = 8.6, p < 0.0001; CS2: t(577) = 3.5, p < 0.0001; CS3: t(576) = 7.26, p < 0.0001). Actions during CS1 declined at all intensities (low: t(1599) = 6.07, p < 0.0001; medium: t(1599) = 5.43, p < 0.0001; high: t(1599) = 6.93, p < 0.0001), as did actions during CS3 (low: t(1606.7) = 4.47, p = 0.0012; medium: t(1599) = 4.29, p = 0.0026; high: t(1607) = 6.79, p < 0.0001). Actions during CS2 were reduced at low intensity (t(1611) = 3.4, p = 0.0018), but not at medium (t(1603) = 2.09, p = 0.10) or high intensity (t(1611) = 1.9, p = 0.052). CS2 elicited more actions than CS3 (t(2315) = 7.13, p < 0.0001). Action latencies increased in AA39 relative to AA19 for CS1 (t(1001) = 8.27, p < 0.0001) and CS2 (t(1222) = 7.5, p < 0.0001), but not for CS3 (t(1198) = 1.16, p = 0.68). CS1 latencies increased at medium (t(2160) = 5.51, p < 0.0001) and high intensity (t(2166) = 6.59, p < 0.0001), but not at low intensity (t(2170) = 3.15, p = 0.38). CS2 latencies increased at low (t(2222) = 5.53, p < 0.0001) and medium intensity (t(2210) = 4.34, p = 0.0042), but not at high intensity (t(2171) = 3.68, p = 0.061). **ITCs**. ITC frequency decreased across task phases from noUS to AA19 (t(271) = 2.78, p = 0.0057) and from AA19 to AA39 (t(271) = 3.34, p = 0.0018). During noUS, ITC rates did not differ among CSs. During AA19, ITCs were lower following CS1 than CS2 (t(2315) = 8.4, p < 0.0001) or CS3 (t(2315) = 4.37, p < 0.0001), and CS2 ITCs exceeded CS3 (t(2315) = 4.1, p < 0.0001). A similar pattern was observed in AA39, except that ITCs following CS2 and CS3 did not differ. ***C***, Same data as in B, shown averaged across tone intensities.

We analyzed behavioral and neural data using linear mixed-effects models with the following fixed factors: *Task* (3 levels: noUS, AA19, AA39), *CS* (3 levels: CS1, CS2, CS3), *Tone Intensity* (3 dB levels: low, medium, high) and G*roup* (2 levels: mPFC, VI), as well as nested random effects (sessions nested within mice). Three behavioral variables were measured: actions, action latencies, and ITCs. Actions were defined as crossings between shuttle-box compartments during CS presentations, though their meaning varied by task as described below. For behavioral analyses (Fig. 2), data from the mPFC and VI groups are presented in combination after confirming no Group factor effect in the model (Fig. 2B and 2C show the same data; in 2C values are averaged across tone intensity).

In the noUS task (5 sessions; Fig. 2A), the three CSs occurred in random order during 7-s action windows under an *ignore* contingency (Ig), in which they predicted no outcome but were terminated if the mouse crossed compartments (*spurious action*). Thus, in noUS, crossings were spurious actions to initially neutral cues, providing measures of baseline exploratory responding and CS neutrality. We found that mice showed no difference in the number of spurious actions across CS1–CS3, indicating that the cues were initially neutral (Fig. 2B, open gray squares; Fig. 2C black circles left panel). However, within each CS, tone intensity significantly modulated responding as high-intensity tones elicited more spurious actions than low-intensity tones.

Notably, CS2 at high intensity produced significantly more crossings than CS2 at medium intensity, suggesting that this tone was perceived as more salient. The percentage of actions to CS presentations during noUS (Fig. 2B) provide an estimate of baseline responding in the absence of learning, reflecting crossings that coincide with cue presentation by chance, as well as responses driven by cue physical saliency. Once US contingencies are introduced and associated with particular cues, this baseline shifts toward more selective responding, as indicated by reductions in responding to non-associated cues and in ITCs per trial described next.

In the AA19 task (7 sessions; Fig. 2A), stimulus presentation was identical to noUS, except that CS1 now incorporated an *active avoidance contingency* (AA), in which it predicted an aversive unconditioned stimulus (US) at the end of the 7-s window unless the mouse crossed compartments (*active avoid*). Failure to avoid resulted in rapid escape responses triggered by the US (*AA error*). Thus, crossings during CS1 were correct active avoids, whereas failures to cross were active avoid errors (escapes). CS2 and CS3 remained non-predictive, as in noUS sessions. We found that in AA19 (Fig. 2B, red circles) mice made significantly more actions during CS1 compared to the noUS phase (Fig. 2C, black circles middle panel), indicating acquisition of active avoidance behavior to prevent the US. This increase was observed for all CS1 intensities. Spurious actions during CS2 and CS3 remained like noUS levels at low and medium intensities but rose significantly at high intensity, suggesting that discrimination between tones became more difficult as loudness increased. Within AA19, the number of active avoids during CS1 was markedly higher than spurious actions during CS2 or CS3, while CS2 and CS3 did not differ. Together, these results confirm robust and selective acquisition of active avoidance responding to CS1 and demonstrate that high intensity non-predictive cues generate discrimination errors.

In the AA39 task (7 sessions; Fig. 2A), CS1 was unchanged, continuing to signal that the animal must act to avoid the US, while CS2 now carried a *passive avoidance contingency* (PA); crossing during CS2 resulted in a brief (0.5 s) US (*PA error*), which mice learned to avoid by withholding the action. CS3 remained unpredictive. Thus, crossings during CS2 were passive avoid errors, while withholding the action constituted a proper passive avoid. We found that in AA39 (Fig. 2B, cyan triangles), as in AA19, mice made significantly more actions during CS1 than during CS2 or CS3 (Fig. 2C black circles right panel). Compared to AA19, overall responding decreased across all CSs, indicating a general suppression of action when the PA contingency was introduced, consistent with increased task difficulty. Active avoids to CS1 declined at all tone intensities, as did spurious actions to CS3. In contrast, CS2 actions—representing passive avoid errors in AA39—were selectively reduced at the lower intensities; mice successfully postponed actions to CS2 at low and medium intensities (<23% passive avoid errors), but at the high intensity they failed to withhold action on >50% of trials. Consequently, CS2 elicited more passive avoid errors than CS3 elicited spurious actions, driven largely by the high-error rate to CS2-high. Together, these results indicate that introducing the passive avoidance contingency reduced overall action rates but that high-intensity tones impaired response inhibition during CS2 trials likely because the loudness makes discrimination vs CS1 challenging.

We next analyzed action latencies, which can reflect increased caution or decision hesitation in challenging environments (Zhou et al., 2022). Action latencies did not differ between the noUS and AA19 phases for any CS (Fig. 2C). During AA39, however, latencies increased relative to AA19 for both CS1 active avoids and CS2 passive avoid errors, but not for CS3 spurious actions. When examined by tone intensity (Fig. 2B), CS1 avoidance latencies in AA39 (relative to AA19) increased at medium and high intensities, but not at low intensity. The absence of an effect at low intensity likely reflects already prolonged latencies in AA19, consistent with reduced detectability of low-salience tones. Latencies of CS2 actions—representing passive avoid errors in AA39—also increased, particularly at low and medium intensities. These elevated latencies suggest that mice were more cautious hesitating longer before initiating actions under more challenging conditions that required discrimination, consistent with greater cognitive demand and response opposition during AA39.

Mice exhibited generally low ITC rates across all phases, indicating that CS-evoked actions occurred against a backdrop of minimal crossings outside of CS windows. Although ITCs were never punished (Zhou et al., 2022), their frequency decreased progressively as contingencies were added (Fig. 2B,C) reflecting reduced exploratory behavior as the task context became more demanding.

### Movement dynamics during contingency changes

We next examined how movement dynamics evolved across task phases. Movement was quantified in three window epochs relative to CS onset: *baseline* (−0.5 to 0 s pre-CS), *orienting* (0–0.5 s post-CS), and *action* (0.5–7 s post-CS). In addition, a *from-action* window (−2 to 2 s relative to door crossing) captured movement surrounding the action event, defined as the moment the animal crossed the compartment door. Movement and neural measures in the three response windows (orienting, action, and from-action) were baseline-subtracted at the trial level, using the −0.5 to 0 s pre-CS baseline window. In this section, we provide an overview of movement across task phases. Tone intensity was included as a factor in all statistical models.

However, for clarity of presentation, intensity is sometimes collapsed in figures to reduce dimensionality and emphasize the other task-related factors. This reflects its role of increasing behavioral variability (error rates), thereby improving sensitivity for estimating the effects of the other task variables.

Movement was analyzed using the same linear mixed-effects framework as the behavioral analyses, with the addition of an *Outcome* factor (2 levels: Action, No-Action) indicating whether a crossing occurred during the CS interval. The Action vs No-Action contrasts in the from-action window were contingency-specific. During the AA contingency, CS1 compared active avoids (Action) with escapes (No-Action) aligned to the crossings. During the PA contingency, CS2 compared passive avoid error crossings (Action) with the equivalent action epoch of correct passive avoid trials (No-Action), which contain no crossings. Similarly, during the Ig contingency, CS1-3 compared spurious crossings (Action) with the equivalent action epoch of ignored trials (No-Action), which contain no crossings. Figure 2-Supplement 1A,B,C show baseline-corrected speed traces aligned to CS onset for Action (A) and No-Action (B) trials and aligned from-action (C). Fig. 2-Supplement 2A,B shows model-derived marginal means of movement averaged across tone intensities (for simplicity). However, the model includes tone intensity, and its effects are described in the supplement legend.

Baseline movement varied systematically across task phases and actions (Fig. 2-Supplement 2A,B Baseline). There were strong main effects of Task and Outcome, as well as significant interactions. During noUS, spurious actions were preceded by higher baseline speed than No-Action trials, indicating that crossings during neutral cues occurred preferentially during periods of elevated motor activity prior to CS onset. In AA19, this pattern persisted for spurious actions during CS2 and CS3, but baseline speed no longer differed between active avoids and escapes (No-Action) during CS1. This indicates that cued active avoids were stimulus-driven rather than determined by pre-existing motor state. In AA39, elevated baseline movement continued to precede spurious actions and was also evident prior to CS2 passive avoid errors and, to a lesser extent, CS1 active avoids, consistent with degraded performance under increased task demands. Across tasks, baseline movement decreased from noUS to AA19, but did not decrease further in AA39, suggesting that baseline motor activity stabilized once the first avoidance contingency was learned.

We next examined CS-evoked orienting responses (Fig. 2-Supplement 2A,B Orient), defined as rapid head movement immediately following CS onset (Fig. 2-Supplement 1A,B), which are known to vary with both tone intensity and task-related behavioral state (Zhou et al., 2023). Orienting magnitude was strongly dependent on tone intensity and was modulated by task phase in a CS-specific manner. These effects were most pronounced at high intensity, where orienting responses to CS1 and CS3 increased from noUS to AA19, whereas CS2 showed comparatively weaker modulation. Because orienting responses could confound subsequent movement measures, the action window was defined to exclude this epoch, allowing isolation of movement associated with action execution.

Movement during the action window or aligned from-action showed similar patterns that were strongly influenced by CS and tone intensity, with other higher-order interactions (Fig. 2–Supplement 1C; Fig. 2–Supplement 2A,B Action). The effect of each CS on movement differed across task phases, and these differences depended on cue intensity. As expected, movement was also strongly influenced by whether an action occurred, with the magnitude of action-related movement effect varying across CSs, task phases, and tone intensities.

Comparing Action (or No-Action) trials between tasks is useful for isolating contingency effects. Focusing on Action trials in the action window, CS1 active avoids during AA19 elicited greater movement than spurious actions during noUS across all intensities (Fig. 2-Supplement 1A; Fig. 2-Supplement 2A,B From Action), consistent with active avoidance learning. Smaller increases in movement were also observed for spurious actions to CS2 and CS3 when comparing noUS and AA19, but these effects were limited to medium and high intensities, likely reflecting discrimination errors relative to CS1. Within AA19, CS1 active avoids also elicited greater movement than CS2 or CS3 spurious actions, whereas in noUS, movement did not differ among CSs when tone intensities were pooled. When examined by intensity, CS2 at high intensity evoked greater movement than other high-intensity cues during noUS, and in AA19 no longer differed from CS1 at high intensity, consistent with elevated salience and reduced discriminability from CS1.

Extending this analysis to the more complex AA39 task revealed contingency-specific changes in movement dynamics. Movement during CS2 passive avoid errors increased in AA39 relative to CS2 spurious actions in AA19, particularly at medium and high intensities. In contrast, movement accompanying CS1 active avoids was largely unchanged between AA19 and AA39, whereas movement during CS3 spurious actions decreased in AA39 relative to AA19, primarily at high intensity. Within AA39, CS2 passive avoid errors were associated with greater movement than CS1 active avoids, again most pronounced at medium and high intensities, consistent with movement driven by the punishing consequences of errors during CS2. These CS-aligned effects were similarly observed in analyses aligned from-action (Fig. 2-Supplement 1C; Fig. 2-Supplement 2A,B).

Finally, we examined No-Action trials in AA39 to isolate the effects of the passive avoidance contingency. Movement during correct CS2 passive avoids was reduced relative to ignored (No-Action) trials during CS2 in AA19. A comparable reduction was also observed for CS1 No-Action trials, whereas movement during CS3 No-Action trials was unchanged. This selective suppression of movement during No-Action trials for cues associated with avoidance contingencies suggests that CS1 and CS2 became behaviorally linked in AA39, where withholding action to CS2 and acting to CS1 represented opposing adaptive strategies.

Together, these results show that movement dynamics systematically track task phase, cue identity, tone intensity, and action outcome. Learned active and passive avoidance contingencies were associated with distinct movement signatures, whereas spurious actions during neutral cues primarily reflected baseline motor activity. Increasing task difficulty in AA39 impaired performance, reflecting reduced discriminability at high intensities and competing motivational drives. This task series provides a framework for probing neural dynamics underlying stimulus-driven decision-making and motor output in the context of learned aversive outcomes.

### Fiber photometry ΔF/F model details

We next asked whether GABAergic neurons in mPFC encode behavioral contingencies across the avoidance task series, and how these signals compare to those in VI. Fiber photometry ΔF/F signals from both regions were analyzed using linear mixed-effects models analogous to those used for behavior and movement (Fig. 2), with ΔF/F as the dependent variable and Group (mPFC, VI) as an additional fixed effect (Fig. 3). The same time windows defined for movement analyses (baseline, orienting, action, and from-action) were also used for the neural analyses.

**Figure 3.**
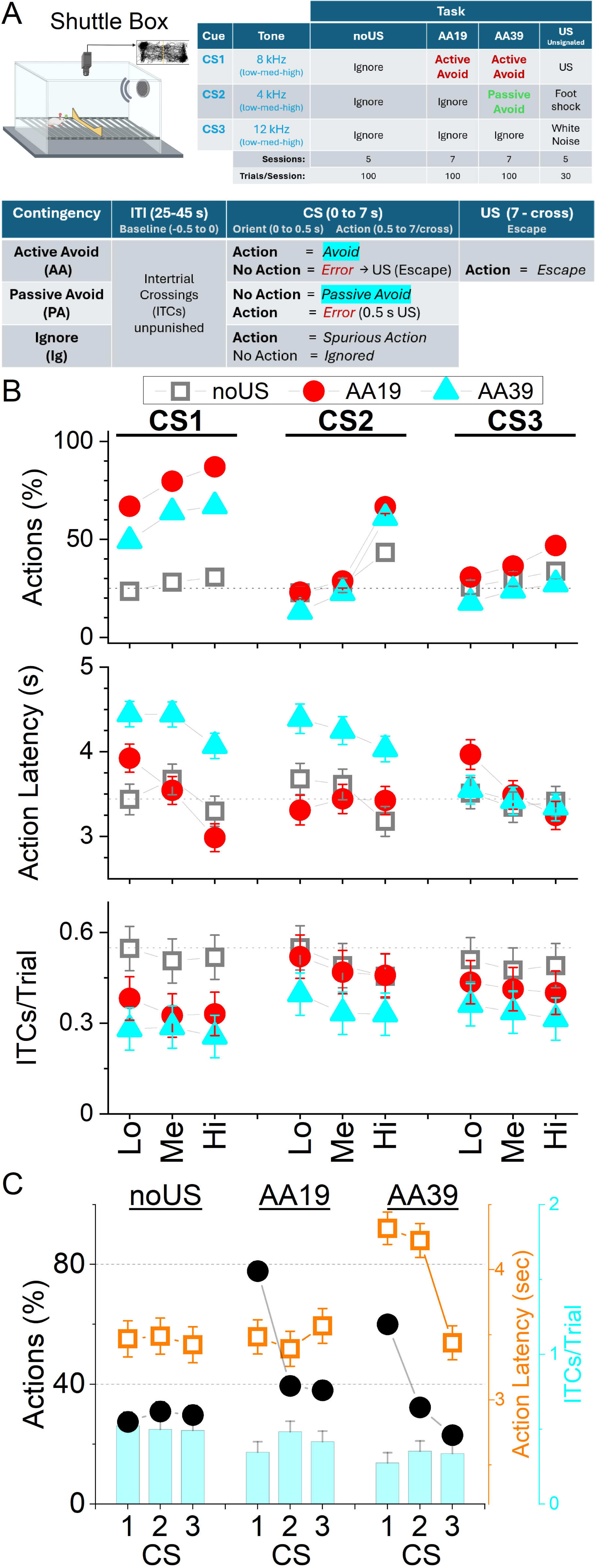
Simple (AA19) and challenging (AA39) avoidance contingencies mixed-effects models of ΔF/F activity controlling for movement and baseline activity. Population estimates of marginal means from ΔF/F fiber-photometry mixed-effects models for AA19 (***A****, **B***) and AA39 (***C****, **D***) tasks, shown separately for mPFC neurons (***A****, **C***) and VI neurons (***B****, **D***). Each panel set contains four rows corresponding to the baseline, orienting, action (CS-aligned), and from-action model windows. Within each row, the left and middle panels compare contingencies (Ignore, Active Avoid, Passive Avoid; Ig, AA, PA) for Action and No-Action trials, and right panel compare Action vs. No-Action outcomes. Actions not classified as AA, PA, Escape, or PA Error are spurious actions. Transparency in the Action window indicates that movement was not controlled between outcomes in that model. ***A****, **B***, AA19 models included Contingency (Ig, AA), Outcome, Group, and covariates. **Baseline window**. A significant Contingency × Outcome interaction was observed (χ²(1) = 12.82, p = 0.00034). No Contingency × Group or three-way interaction was detected. (R^2^_m_ = 0.22; R^2^_c_ = 0.26) **Orienting window**. A significant Outcome × Group interaction was detected (χ²(1) = 5.25, p = 0.022). (R^2^_m_ = 0.2; R^2^_c_ = 0.36) **Action window**. Two-way interactions were present among factors, with a significant Contingency × Group interaction (χ²(1) = 9.25, p = 0.0023). No three-way interaction was detected. Spurious actions during Ig trials did not differ from active avoids during AA trials in either region. (R^2^_m_ = 0.53; R^2^_c_ = 0.66) **From-Action window**. (From-Action is defined as the time at which the animal crosses the door) A significant Contingency × Outcome × Group interaction was observed (χ²(1) = 9.56, p = 0.0019). No differences were detected between spurious actions and active avoids in either region. (R^2^_m_ = 0.42; R^2^_c_ = 0.6) ***C****, **D***, AA39 models included Contingency (Ig, AA, PA), Outcome, Group, and covariates. **Baseline window**. A significant Contingency × Outcome interaction was detected (χ²(2) = 20.77, p < 0.0001). No interaction with Group was observed. (R^2^_m_ = 0.23; R^2^_c_ = 0.31) **Orienting window**. Significant Contingency × Outcome (χ²(2) = 24.87, p < 0.0001) and Contingency × Group interactions were detected (χ²(2) = 31.58, p < 0.0001), with no three-way interaction. (R^2^_m_ = 0.3; R^2^_c_ = 0.47) **Action window**. Significant two-way interactions were observed, including a Contingency × Group interaction (χ²(2) = 24.14, p < 0.0001), as well as a significant Contingency × Outcome × Group interaction (χ²(2) = 6.35, p = 0.041). Spurious actions during Ig trials did not differ from active avoids in either region. Correct passive avoids differed from correct ignores in mPFC (t(1301) = 4.14, p < 0.0001). (model fit: R^2^_m_ = 0.58; R^2^_c_ = 0.72) **From-Action window**. A significant Contingency × Outcome × Group interaction was detected (χ²(2) = 11.78, p = 0.0027). Active avoids differed from spurious actions in mPFC (t(1362) = 2.52, p = 0.011) but not in VI. (model fit: R^2^_m_ = 0.5; R^2^_c_ = 0.67) ***E***, Mean ΔF/F (top) and movement (bottom) traces for mPFC and VI neurons during the unsignaled escape task, showing interleaved trials of footshock alone, white noise (WN) alone, and the full US, aligned to US onset. ***F***, Population estimates of marginal means from ΔF/F mixed-effects models (including movement covariates) during the escape window (0-4 s from US onset). In the **baseline window**, a strong effect of baseline movement was observed (χ²(1) = 14.31, p = 0.00015), with no main effects or interactions involving US or Group. During the **escape window**, significant effects of escape movement (χ²(1) = 12.09, p = 0.00052), baseline movement (χ²(1) = 4.15, p = 0.041), and baseline ΔF/F (χ²(1) = 140.14, p < 0.0001) were detected. After accounting for covariates, a significant main effect of US (χ²(2) = 33.02, p < 0.0001) and a US × Group interaction were observed (χ²(2) = 11.38, p = 0.0034). In mPFC, footshock alone differed from the full US (t(105) = 3.89, p = 0.00034) and white noise alone (t(105) = 6.13, p < 0.0001); no corresponding contrasts were significant in VI.

Across task phases, both mPFC and VI exhibited robust CS- and task-related ΔF/F modulation (Fig. 3-Supplement 1 shows ΔF/F and associated movement traces). Much of this activity was temporally aligned to orienting responses at CS onset and to subsequent avoid or escape movements, with faster movements generally associated with larger ΔF/F transients. These observations motivated explicit control for movement and baseline activity in the models (Fig. 3 and Fig. 3-Supplement 2).

To dissociate contingency-related neural modulation from movement-related signals, all models included covariates for window-specific head speed, baseline head speed, and baseline ΔF/F (the latter excluded from the baseline window). Covariates were z-scored within each window so that estimated marginal means reflected ΔF/F at average covariate values. In the action window, movement was z-scored separately for Action and No-Action trials, because these outcomes differ substantially in movement magnitude; as a result, this model (window) controls for speed within, but not between, outcomes. In contrast, the from-action window explicitly enables comparison across actions while controlling for movement. Random effects consisted of sessions nested within mice (the behavioral data for these animals was shown in Fig. 2).

Two complementary modeling approaches were used. First, task-specific models were fit separately for AA19 and AA39 to directly compare contingencies and outcomes within each task, without influence from other phases (Fig. 3). Second, a *full model* incorporating all task phases (noUS, AA19, AA39) was used to assess task-dependent changes in neural activity (Fig. 3–Supplement 2).

### mPFC GABAergic activity preferentially encodes error outcomes

We first summarize the main effects and interactions identified by the full model, with emphasis on how contingency encoding changes across task phases and cortical regions (Fig. 3-Supplement 2). Then, we focus on the contingencies within the AA19 and AA39 tasks models (Fig. 3). In every case, we describe the baseline, orienting, action, and from-action window models in sequence.

#### Full model

In all the windows, the covariates had strong effects on ΔF/F, indicating the importance of including them in the models (Fig. 3-Supplement 2). After controlling for the covariates, baseline ΔF/F activity varied with task contingencies and outcomes, but this was largely similar across cortical regions, indicating a shared modulation of context-related excitability (Fig. 3-Supplement 2 Baseline). Similarly, during the orienting window, both regions showed weak contingency-related modulation, with mPFC exhibiting slightly greater sensitivity to task transitions (Fig. 3-Supplement 2 Orient).

In the action window, both mPFC and VI enhanced their activation during passive avoid errors in AA39 compared to spurious actions in AA19 (Fig. 3-Supplement 2 Action). In addition, only mPFC encoded correct passive avoids in AA39 as they produced stronger activation than No-Action trials in AA19, but this CS2 change also occurred between noUS and AA19 and may be contextual—unrelated to the PA contingency in AA39. Active avoids in AA19 were associated with enhanced activation vs spurious actions in noUS, but this may reflect other factors than avoidance acquisition since spurious actions showed similar effects across these tasks, suggesting a contextual-related effect.

During the from-action window, mPFC neurons exhibited stronger activation to escapes than active avoids and to passive avoid errors than spurious actions across tasks, whereas these effects were weaker or largely absent in VI (Fig. 3-Supplement 2 From Action). Most prominently, there were no differences between active avoids in AA19 and spurious actions in noUS, highlighting that acquisition of active avoidance was not encoded around the time of action.

After controlling for baseline and movement-related factors, these results reveal some regional functional dissociation in which mPFC encodes avoidance errors around the action. However, because avoidance errors are followed by aversive stimulation, neural activity in the from-action window may reflect a combination of movement-related and punishment-related signals.

#### AA19 simple avoidance context model

The AA19 model (Fig. 3A,B) focuses on the simple avoidance context. These models combine the CS trials by their contingencies (Ig, AA, PA) and outcomes (Action, No-Action). In AA19, baseline activity predicted behavioral outcomes differently across contingencies, but this relationship did not differ between regions. In mPFC, higher baseline ΔF/F during Ig trials preceded spurious actions, whereas in VI higher baseline activity during AA trials preceded escapes (Fig. 3A,B Baseline).

During the orienting window, outcome-dependent modulation differed between regions (Fig. 3A,B). mPFC showed elevated activity during spurious actions, whereas VI showed minimal modulation (Fig. 3A,B Orient). During the action window, escapes produced stronger activation than correct ignores in mPFC, but spurious actions did not differ from active avoids in either region, indicating no encoding of active avoidance (Fig. 3A,B Action).

In the from-action window, mPFC—but not VI—showed selectively elevated activation during escapes relative to active avoids, demonstrating region-specific encoding of failed active avoidance (Fig. 3A,B From Action). Importantly, neither region differentiated active avoids from spurious actions, confirming that mPFC does not encode active avoidance in this simple context.

#### AA39 difficult avoidance context model

The AA39 model (Fig. 3C,D) focuses on the more challenging context where mice must discriminate tones to select opposing actions. In AA39, baseline ΔF/F again predicted outcome differently across contingencies, but similarly in both regions; higher baseline activation preceding Ig trials was associated with spurious actions in both regions (Fig. 3C,D Baseline). Spurious actions showed higher baseline activation than active avoids in both mPFC and VI, and compared to passive avoid errors in mPFC.

During the orienting window, contingency-related modulation was stronger than in AA19 (Fig. 3C,D Orient). Passive avoid errors were associated with higher orienting activation than correct passive avoids in both regions, while escapes had stronger orienting responses than active avoids selectively in mPFC. In VI, spurious actions during Ig trials elicited stronger orienting activation than correct ignores.

In the action window, passive avoid errors produced stronger activation than active avoids and spurious actions in both regions (Fig. 3C,D Action). Notably, correct passive avoids elicited greater activation than correct ignores in mPFC, despite both involving action withholding, indicating encoding of deliberate inhibition rather than absence of action. Like in AA19, active avoids did not differ from spurious actions, again indicating no encoding of active avoidance from CS onset.

In the from-action window, mPFC showed robust enhancement during escapes and passive avoid errors relative to successful outcomes, whereas VI showed minimal modulation (Fig. 3C,D From Action). This effect may reflect higher sensitivity to the aversive US outcome in mPFC. Unlike AA19, active avoids in AA39 elicited stronger activation than spurious actions in mPFC but not VI, indicating that mPFC encodes active avoidance only around the time of action under challenging conditions.

#### Summary

Together, these results indicate two main findings. First, mPFC does not encode active avoidance when it emerges in simple contexts where decisions are easy and errors rare. In contrast, in challenging contexts, where cautious responses emerge, and failures are frequent, mPFC encodes passive avoidance and somewhat active avoidance, but only around the time of the action. Because mPFC activity increased during both active and passive avoidance, this activation likely reflects elevated cognitive demands rather than encoding of the specific action selected. The second finding is that error-related activation linked to punishment was present in both mPFC and VI but appeared more strongly and consistently in mPFC across contingencies, suggesting that GABAergic neurons in this region are particularly sensitive to aversive outcomes.

### Error encoding in mPFC reflects sensitivity to unsignaled aversive stimulation

To further probe how aversive outcomes are represented in mPFC and VI GABAergic neurons, we examined neural activity during escape-only sessions in which the unconditioned stimulus (US) was delivered without predictive cues (n=7 mice in mPFC, n=5 mice in VI; Fig. 3E,F). In these sessions, mice experienced randomly interleaved presentations of footshock alone, white noise alone, or the full US composed of white noise followed by footshock (same US used in the tasks). We applied mixed-effects models with the same covariates as during the other tasks to the baseline and escape (0-4 s from US onset) windows.

After controlling for the covariates, baseline ΔF/F activity did not differ across stimulus types or between brain regions, indicating that pre-stimulus neural activity was comparable across conditions. In contrast, during the escape window following US onset, neural responses diverged across stimulus types in a region-specific manner (Fig. 3E,F Action). In mPFC, footshock presented in isolation elicited stronger activation than either white noise alone or the full US, despite the latter containing the same aversive footshock component. This enhancement was specific to mPFC and was not observed in VI neurons, which showed similar activation levels across all stimulus conditions.

The selective mPFC enhancement to footshock alone suggests that mPFC GABAergic neurons are particularly sensitive to aversive outcomes when they occur without predictive sensory signals. In contrast, when the footshock was accompanied by an auditory white noise cue as part of the full US, mPFC activation was attenuated, consistent with reduced neural engagement when aversive events are signaled. The absence of this modulation in VI indicates that a sensitivity to unsignaled punishment is not a general property of cortical GABAergic activity but rather reflects a specialized role of mPFC in encoding aversive events, perhaps under conditions of uncertainty or unpredictability.

### Classification of GABAergic mPFC neurons by their relationship to movement

We next used miniscopes to image activity from individual GABAergic mPFC neurons (n=2217 neurons from 5 mice) while mice performed AA19 (Fig. 4; n=367) and AA39 (Fig. 4-Supplement 1; n=1850). Rather than classifying neurons based on stimulus-locked responses alone, we leveraged the prominent movement sensitivity observed in the population photometry data to organize neurons according to their relationship with movement. This approach allowed us to test whether movement sensitivity defines functional subgroups that differ in their encoding of task contingencies. Classification was performed in two steps. First, we computed the cross-correlation between head speed and ΔF/F for each neuron. Second, we applied k-means clustering to these cross-correlation functions to group neurons based on the timing (lag) and strength of the relationship between neural activity and movement. This analysis identified five neuron classes (Fig. 4 and Fig. 4-Supplement 1, inset “a”). One class showed strong correlations centered near zero lag (Class 5, cyan), three classes exhibited progressively weaker correlations with increasing positive lag (Class 4, blue; Class 2, red; Class 1, black), consistent with movement preceding neural activation to varying degrees, and one class showed negative correlations with movement (Class 3, green), indicating suppression during movement. The neuron Class was included as a fixed effect factor in the mixed-effects models, with neuron identity nested within mouse as a random effect. Equivalence testing confirmed that movement measures overlapped across neuron classes (Fig. 4 and Fig. 4-Supplement 1 speed traces), indicating that class-related effects could not be attributed to systematic differences in movement across task contingencies.

**Figure 4.**
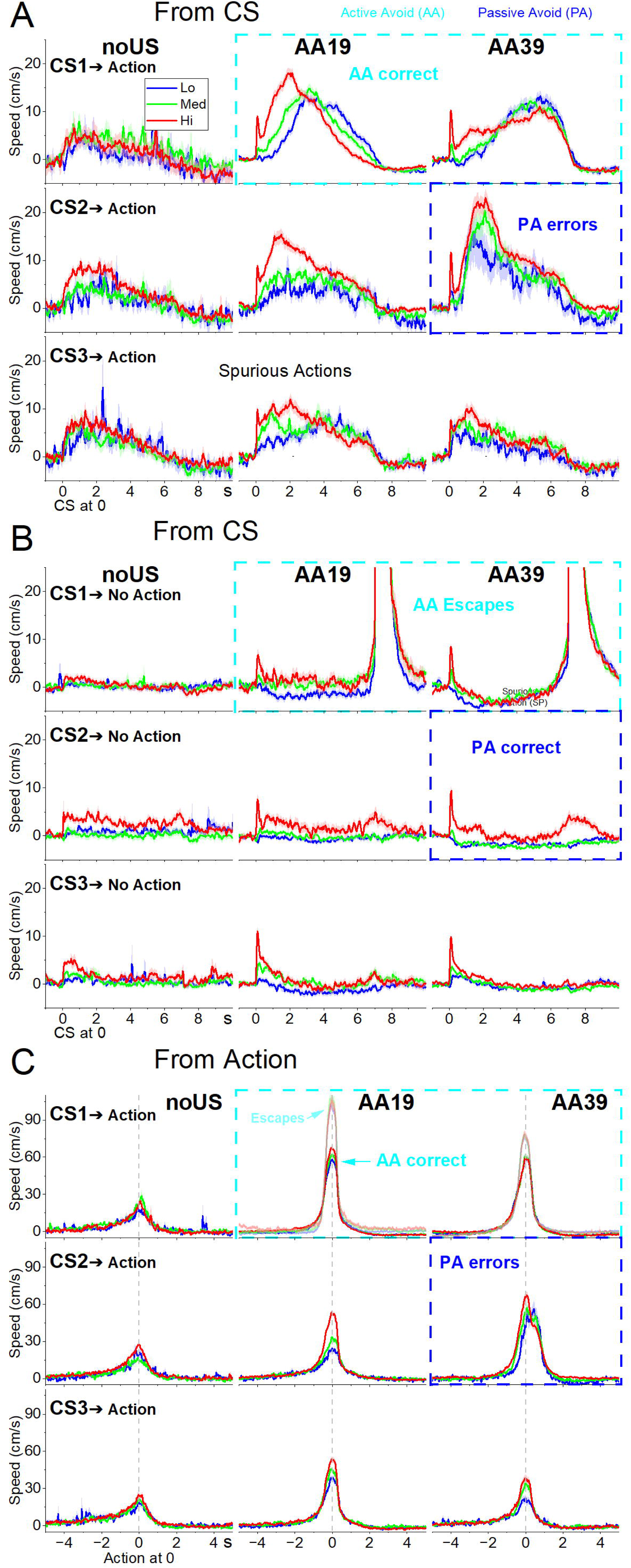
Movement-related neuron classes during a simple avoidance context (AA19 task). ***A***, Mean ΔF/F traces (top) and corresponding movement traces (bottom) for five mPFC GABAergic neuron classes identified by k-means clustering of the cross-correlation functions between speed and ΔF/F (inset **a**). Class 1 (black) and Class 2 (red) exhibited moderate positive correlations with movement, Class 3 (green) showed negative correlations, and Classes 4 (blue) and 5 (cyan) exhibited strong positive correlations, with Class 5 centered near zero lag. Traces are aligned to CS onset, for CS1–CS3, separated by contingency (Ignore, Ig; Active Avoid, AA) and behavioral outcome (action vs No-Action), aligned to CS onset. In AA19, CS1 signals active avoidance and an Action outcome is the correct response. ***B***, Same as A, aligned from-action (defined as the time at which the animal crosses the door) for Action outcomes (no traces exist from-action for No-Action trials). ***C,D***, Estimated marginal means of ΔF/F from linear mixed-effects models for the baseline, orienting, action, and from-action windows. Models compared contingencies (Ig vs AA) and outcomes (action vs No-Action) across neuron classes. ***C***, contrasts across contingencies within outcomes. ***D***, contrasts between outcomes within each contingency and neuron class. Transparency in the action window indicates that movement was not controlled between outcomes in that model. **Baseline window**. The model revealed a significant Contingency × Outcome interaction (χ²(1) = 44.01, p < 0.0001) and a significant Contingency × Outcome × Class interaction (χ²(4) = 11.21, p = 0.024), indicating class-dependent differences in pre-CS activity across contingencies and outcomes. In Classes 1 and 2, baseline ΔF/F differed between correct active avoids and spurious actions, while in Classes 1, 2, and 4, baseline activity differed between escapes and correctly ignored CSs. (R^2^_m_ = 0.09; R^2^_c_ = 0.14) **Orienting window**. There was a significant main effect of neuron Class (χ²(4) = 124.33, p < 0.0001), indicating differences in orienting-related ΔF/F across classes. The Contingency × Outcome interaction with Class was weak (p = 0.08). In Class 3, orienting-related activity differed between active avoids and spurious actions. In Ig trials, orienting-related activity differed between spurious actions and correctly ignored CSs in Classes 2, 3, and 5. (R^2^_m_ = 0.17; R^2^_c_ = 0.39) **Action window**. The model revealed a significant Contingency × Outcome interaction (χ²(1) = 20.47, p < 0.0001), with no significant three-way interaction involving Class (p = 0.08). Across classes, ΔF/F differed between escapes and correctly ignored CSs, except in Class 1. Differences between active avoids and spurious actions were class dependent, with opposite patterns observed in Classes 2 and 3. Comparisons between Action and No-Action outcomes are shown for completeness but are not controlled for movement in this window. (R^2^_m_ = 0.35; R^2^_c_ = 0.51) **From-Action window**. A significant Contingency × Outcome × Class interaction was observed (χ²(4) = 17.33, p = 0.0016), indicating class-specific differences in action-aligned activity. Only Class 5 showed a difference between active avoids and spurious actions. In most other classes, ΔF/F differed between escapes and active avoids. (R^2^_m_ = 0.29; R^2^_c_ = 0.47)

### Movement-related neuron classes during a simple avoidance context

Figure 4A,B show the activity of these neuron classes and associated movement during AA19, aligned to CS onset (A) and to action occurrence (B) for each contingency (Ig, AA) and behavioral outcome. Class 5 neurons (cyan, 6.8%), which exhibited near-zero-lag correlations with movement, showed the strongest activation, followed by Class 4 neurons (blue, 16.9%), whose activity closely tracked movement. Class 3 neurons (green, 15.8%) were inhibited around the time of action, whereas Class 1 (black, 27.3%) and Class 2 (red, 33.2%) neurons showed relatively weak activation. Notably, Class 5 neurons also exhibited elevated activity during No-Action trials across CS1, CS2, and CS3, when movement remained near baseline. These responses were most prominent at CS onset, indicating that this class may be sensitive to cue presentation independent of overt movement. How these neuron classes differ in contingency-and outcome-dependent encoding is examined in the following analyses.

In the mixed-effects models (Fig. 4C,D), covariates related to movement and baseline activity strongly influenced ΔF/F across all windows, emphasizing the importance of controlling for these factors. During the baseline window, pre-CS activity differed across neuron classes depending on contingency and outcome. In particular, Classes 1 and 2 showed lower baseline activity preceding correct active avoids than spurious actions, whereas escapes were preceded by elevated baseline activity relative to correctly ignored cues in Classes 1, 2, and 4 (Fig. 4C Baseline). Thus, across these three classes, elevated baseline activity was associated with active avoidance errors rather than successful avoidance (Fig. 4D Baseline).

During the orienting window (Fig. 4C,D Orient), activity patterns were dominated by differences between neuron classes rather than by contingency or outcome. Only Class 3 neurons showed higher activation during active avoids compared to spurious actions, indicating selective encoding of avoidance during the orienting response (Fig. 4C Orient). In Ig trials, orienting-related activity during spurious actions exceeded that during correctly ignored cues in Classes 2 and 5, whereas Class 3 neurons showed the opposite pattern. This inversion is consistent with the negative correlation between Class 3 activity and movement.

During the action window (Fig. 4C,D Action), outcome-related differences were observed across neuron classes. AA errors leading to escapes elicited stronger activation than correctly ignored cues in most classes except Class1, but this may reflect late initiated avoid attempts that failed to reach the door before the end of the action epoch. However, selective encoding of active avoidance was again restricted to Class 3 neurons, which showed higher activation during active avoids than spurious actions. In contrast, Class 2 neurons exhibited the opposite pattern. Thus, as in the orienting window, only Class 3 neurons selectively encoded active avoidance with higher activation relative to spurious actions (Fig. 4C Action). Comparisons between Action and No-Action outcomes (Fig. 4D Action) are deemphasized in this window because movement was not controlled across outcomes by design.

In the from-action window (Fig. 4C,D From Action), action-related activity depended strongly on neuron class. Only Class 5 neurons showed stronger activation for active avoids than for spurious actions, indicating that this subset of GABAergic neurons—characterized by strong, near-zero-lag correlations with movement—selectively encodes the avoidance action itself when aligned to action execution. In contrast, most other classes showed stronger activation during escapes than active avoids, consistent with responses to aversive stimulation. This escape-related enhancement was absent or attenuated in Class 3 and Class 5 neurons, which preferentially encode avoidance from CS onset or the avoidance action, respectively.

Together, these results indicate that during AA19, two neuron classes are linked to active avoidance behavior but in distinct ways. Class 3 neurons, which are inhibited by movement, show weak cue-evoked activation that nonetheless differentiates successful avoidance from spurious actions. Class 5 neurons, which are robustly activated by both movement and cues, selectively encode active avoidance only when activity is aligned to the action itself. The remaining neuron classes primarily reflect responses associated with aversive stimulation rather than avoidance per se.

### Movement-related neuron classes during a difficult avoidance context

We next applied the same analyses to neurons recorded during AA39, a more demanding task that incorporates both active and passive avoidance contingencies. Although recordings were obtained from the same animals as in AA19, the neurons analyzed here represent an independent sample. As before, cross-correlation and clustering analyses identified five neuron classes with similar relationships to movement (Fig. 4–Supplement 1, “a” inset). Figure 4–Supplement 1A,B shows activity from each class aligned to CS onset (A) and from-action (B), together with associated movement traces, separated by contingency and behavioral outcome. In AA39, CS2 signaled the PA contingency, such that actions during CS2 constituted passive avoid errors, whereas withholding action reflected correct passive avoidance performance.

Across classes, Class 5 neurons (Fig. 4-Supplement 1A,B; cyan, 10.9 %), which had peak correlation near zero lag, showed the strongest activation during AA39, followed by Class 4 neurons (blue, 10.3 %), whose activity closely tracked movement. Classes 1 (black, 32.4 %) and 2 (red, 22.7 %) exhibited weaker activation, while Class 3 neurons (green, 23.7 %), whose activity was inhibited in relation to movement, showed minimal activation across conditions. Thus, the overall organization of movement-related neuron classes during AA39 resembled that observed in AA19, despite the added task complexity.

In the mixed-effects models (Fig. 4-Supplement 1C,D), the covariates retained strong effects across all windows. Baseline activity preceding cue onset differed systematically across neuron classes and behavioral outcomes. Classes 1, and 2 showed lower baseline activation for active avoids and passive avoid errors than for spurious actions, while escapes had higher baseline activation than correctly ignored CSs or correct passive avoids in Classes 2, 3, 4, and 5 (Fig. 4-Supplement 1C Baseline). All classes, except Class 3, showed elevated baseline activity preceding spurious actions in Ig trials. In AA trials, escapes were preceded by higher baseline activity in Classes 2, 3, and 4. In PA trials, errors were predicted by higher baseline activity in Class 1 neurons, which had the weakest relation to movement, whereas this relation was inverted for Class 2 neurons. Thus, elevated baseline activation was associated with a greater likelihood of spurious actions or AA errors across most neuron classes, but this relation was more complex in the PA contingency, where Class 1 and 2 neurons showed opposite patterns.

During the orienting window following CS onset, cue-evoked activity depended on both contingency and neuron class. Across most classes, cues predicting AA or PA contingencies elicited stronger orienting-related responses than cues predicting Ig trials, regardless of behavioral outcome (Fig. 4-Supplement 1C Orient). In Classes 1, 5, and to a lesser extent Class 3, AA and PA errors as well as spurious actions were accompanied by heightened CS-evoked activation, suggesting that elevated mPFC activity at cue onset predicts behavioral outcomes.

In the action window, activity was strongly modulated by behavioral outcomes across all neuron classes. For all classes, escapes in AA trials produced stronger activation than correctly ignored Ig trials and correct PA trials, likely related to late avoidance attempts as in AA19. In Classes 2, 4, and 5, PA errors elicited stronger activation than active avoids or spurious actions, consistent with responses to the aversive US. Only Class 5 neurons showed stronger activation for active avoids than for spurious actions, indicating that this subset of neurons—strongly coupled to movement—selectively encodes the avoidance action from CS onset as this effect was also present in the orienting window. Class 5 neurons also encoded passive avoidance, showing stronger activation than during ignored Ig trials, a pattern shared to a lesser extent by Classes 1 and 3 (Fig. 4-Supplement 1C Action). Comparisons between outcomes (Fig. 4-Supplement 1D Action) are deemphasized because movement was not controlled across outcomes by design.

In the from-action window, responses depended strongly on neuron class. Class 5 and 4 neurons showed stronger activation for active avoids than for spurious actions, indicating that these movement-correlated GABAergic neurons encode the avoidance action itself (Fig. 4-Supplement 1C From Action). In these classes, passive avoid errors evoked stronger activation than active avoids, but this activation was comparable to that observed for escapes, consistent with responses driven by the aversive US. In contrast, Class 1 neurons showed the opposite relationship, with passive avoid errors producing less activation than active avoids or spurious actions. Across all classes except Class 1, escapes produced stronger activation than active avoids (Fig. 4-Supplement 1D From Action)., further supporting a general sensitivity to the aversive US across most neuronal classes.

Together, these results indicate that during AA39, the encoding structure of mPFC GABAergic neurons remains broadly like that observed in AA19 but incorporates distinct features consistent with the greater cognitive demands imposed by the added PA contingency. Baseline activity in most classes predicted subsequent errors, suggesting that elevated pre-CS excitability promotes inappropriate or mistimed responses. Class 5 neurons, which in AA19 selectively encoded the avoidance action, now showed strong activation during both correct active and passive avoidance, indicating recruitment for both forms of action control. Their activation also reflected error trials, consistent with heightened sensitivity to aversive stimulation—a feature shared by Classes 1, 2, and 4. In contrast, Class 3 neurons lost the outcome selectivity that distinguished active avoids from spurious actions in AA19, suggesting contextual encoding. Thus, under the more demanding conditions of AA39, Class 5 GABAergic mPFC neurons—those most tightly coupled to movement—encode correct avoidance performance regardless of whether it is expressed through action or inaction.

### Action-related movement defines distinct active avoidance modes

Whereas the previous analysis sorted mPFC neurons based on their correlation with movement, we next asked whether mPFC activity differentially encodes distinct types of active avoidance actions. To capture variation in avoidance behavior, we applied k-means clustering directly to the time series of movement speed aligned to CS onset across all correct active avoidance trials. This approach differs from the preceding one in that it classifies only correct avoidance responses based on their movement dynamics from CS onset (it is not a neuron classification), whereas the previous cross-correlation clustering classified neurons based on their sensitivity to movement across recording sessions.

This analysis revealed three distinct avoidance response modes (Fig. 5-Supplement 1A, bottom gray panel). Mode 1 responses (black) were rapid onset avoids initiated immediately after the orienting movement, which was larger than in the other modes. Mode 2 (red) and Mode 3 (cyan) responses were initiated later and exhibited significantly higher movement speed than Mode 1 when measured from-action, consistent with more cautious, delayed but vigorous avoidance behavior (Zhou et al., 2022).

We next used mixed-effects models to test how mPFC GABAergic neuron activity varied across avoidance modes by considering the ΔF/F time series across recorded neurons (Fig. 5-Supplement 1A, top panel, aligned to CS onset; Fig. 5-Supplement 1B, aligned to avoidance action). In these models (Fig. 5-Supplement 1C), the covariates retained strong effects across all windows. In the baseline and orienting windows, there were no differences across modes, indicating that the selection of avoidance strategy was not influenced by pre-CS mPFC activity. During the action window, both Mode 2 and Mode 3 avoids exhibited higher activation than Mode 1 avoid, and Mode 2 tended to have higher activation than Mode 3 avoids. These differences were stronger in the from-action window (Fig. 5-Supplement 1C From Action).

In summary, mPFC activation scales with behavioral caution consisting of delayed but fast avoidance responses (Modes 2 and 3) consistently evoking the strongest mPFC activation, even after controlling for movement and baseline covariates. This indicates that mPFC activity does not simply mirror motor output but instead reflects higher-order aspects of behavioral control. However, because this approach treats mPFC neurons as a single functional population, we next examined whether distinct GABAergic neuron activations occur across avoidance modes.

### Distinct mPFC GABAergic neurons encode cue and action components across avoidance modes

We next asked whether distinct patterns of mPFC GABAergic neuron activity underlie the different active avoidance modes. To this end, we applied k-means clustering to the ΔF/F time series of individual neurons within each avoidance mode (Fig. 5A,B). In this analysis, activation “*Types*” reflect distinct neuronal response profiles within a given avoidance mode; neurons could contribute to more than one mode if they exhibited avoids in multiple modes, but activation types were treated independently across modes. We show the speed traces for each activation type separately (Fig. 5A,B bottom panels), because activation types are composed of neurons that can be recorded in the same or different sessions. Therefore, the corresponding speed averages can potentially differ across types within a mode, but the movements are remarkably similar (overlapping) despite the large differences in neural activation between the types, indicating that avoidance modes capture similar behavioral outputs.

**Figure 5.**
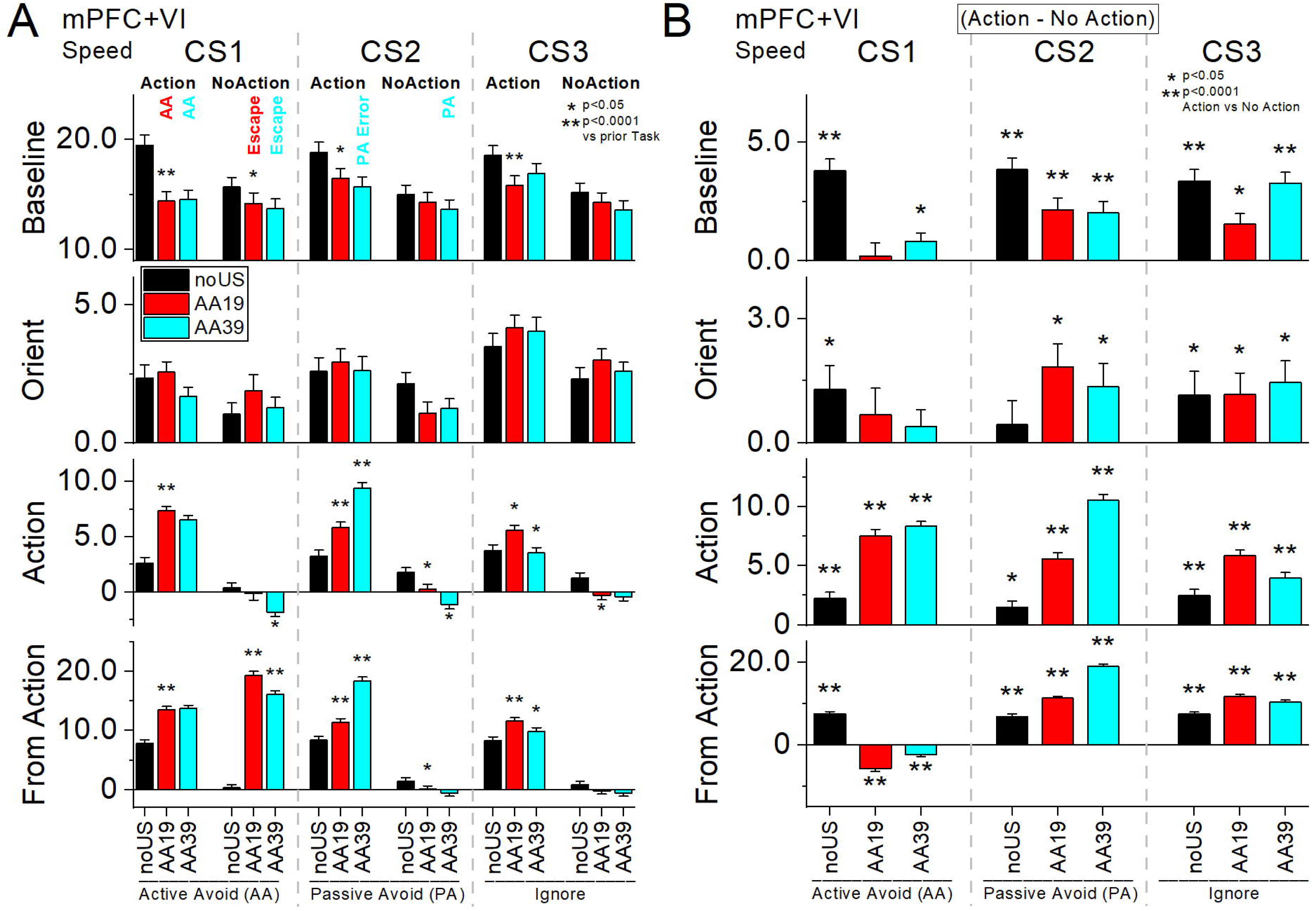
Distinct mPFC GABAergic neuron activation types during active avoidance modes. ***A***, k-means clustering of ΔF/F time series from individual mPFC GABAergic neurons within each active avoidance mode (defined in Fig. 5–Supplement 1) revealed three activation types (a–c). Traces are aligned to CS onset. *Type a* (1–3a) exhibited minimal modulation during avoidance. *Type b* (1–3b) showed a rapid increase in activity at CS onset followed by a gradual decline. *Type c* (1–3c) exhibited selective activation around the time of the avoidance action. Activation types are shown separately for each avoidance mode. *Type b* activation was strongest for Mode 1 and demodulated in Mode 3, while *Type c* activation was more robust in Modes 2 and 3. The bottom panels show the movement (speed) for each of three activation types within each avoid mode. As expected, movement traces overlap across types within a mode, indicating that differences in neural activity are not explained by differences in movement. Minor differences across speed traces arise because some activation types may not be present in all sessions. ***B***, Same as A, aligned from-action (defined as the time at which the animal crossed the door to avoid). ***C***, Population estimates of marginal means (ΔF/F) from linear mixed-effects models for the baseline, orienting, action, and from-action windows, comparing activation Types (a–c) across the three active avoidance modes. **Baseline window**. No differences across activation types or avoidance modes. (R^2^_m_ = 0.09; R^2^_c_ = 0.12) **Orienting window**. Type b activation differed across avoidance modes (Mode 1 vs Mode 3: t(5748) = 2.67, p = 0.02). (R^2^_m_ = 0.12; R^2^_c_ = 0.5) **Action window**. Type a and Type c activation was higher during Mode 2 (Type a: t(4871) = 3.62, p = 0.0008) and Mode 3 (Type a: t(5169) = 3.57, p = 0.0008) compared to Mode 1, whereas Type b showed no mode-dependent differences. (R^2^_m_ = 0.26; R^2^_c_ = 0.71) **From-Action window**. Type b activation was lower during Mode 3 compared to Mode 2 (t(4356) = 4.69, p < 0.0001) and Mode 1 (t(4867) = 2.83, p = 0.0091). Type c activation was higher during Mode 2 (t(4727) = 12.03, p < 0.0001) and Mode 3 (t(4958) = 10.74, p < 0.0001) compared to Mode 1. (R^2^_m_ = 0.15; R^2^_c_ = 0.66)

This analysis revealed three activation types with distinct temporal dynamics within each avoidance mode. *Type a* activation exhibited little modulation during active avoidance (Fig. 5A,B gray traces). *Type b* activation showed a sharp increase in activity at CS onset followed by a gradual decline (Fig. 5A,B green traces), whereas *Type c* activations were selectively activated around the time of the avoidance action (Fig. 5A,B orange traces).

Mixed-effects models revealed that neither activation type showed baseline differences across avoidance modes (Fig. 5C Baseline). During the orienting window, the CS-related *Type b* activation was stronger during the early onset Mode 1 avoids than during the most cautious Mode 3 avoids consistent with encoding of cue or preparatory processes (Fig. 5C Orient). During the action window, the action-related *Type c* activation is larger during the cautious Mode 2 and Mode 3 avoids relative to early onset Mode 1 avoids, whereas *Type b* showed little modulation across modes; *Type a* activations are very small but showed a pattern like *Type c* (Fig. 5C Action). In the from-action window, the CS-related *Type b* activations were reduced in the most cautious Mode 3 avoids compared to the other modes (Fig. 5C From Action). The action-related *Type c* activations were strongly elevated for the cautious Mode 2 and Mode 3 avoids compared to early onset avoid Mode 1.

Together, these results indicate that mPFC GABAergic neurons exhibit complementary activation patterns that map onto distinct epochs of active avoidance. CS-related activation (*Type b*) is greatest for early onset avoids while action-related activation (*Type c*) is strongly elevated during cautious avoids.

### mPFC inactivation has little effect on cued goal-directed behavior under threat

The preceding experiments suggested that mPFC activation may contribute to cued goal-directed behaviors under threat, including both active and passive avoidance. If so, optogenetic inhibition of mPFC should disrupt these behaviors. To test this, we used two complementary optogenetic approaches to inhibit mPFC as mice learned and performed a sequence of tasks: initial active avoidance in AA1; the transition to cautious responding in AA2, where ITCs are punished and avoidance latencies increase; and finally the more demanding AA3, in which mice must discriminate between cues to either generate or postpone an action.

For the first optogenetic inhibition method, we broadly expressed eArch3.0 in mPFC neurons (n = 6) using bilateral AAV-hSyn injections and implanted optical cannulas (Fig. 1A, Fig. 6C). We avoided selectively suppressing GABAergic neurons because this would disinhibit local glutamatergic circuits and produce the opposite of the intended effect—aberrant excitation that would confound behavioral interpretation. In naïve mice, we first asked whether mPFC inhibition impairs acquisition of AA1 when every trial was a CS+Light trial, using green light at three powers (10–30 mW) applied during the avoidance/escape interval. Catch trials without CS or light (NoCS) and ITCs showed typical spurious actions. Mice acquired AA1 at the same rate as controls, with no detectable impairment (Fig. 6A, black symbols). Under the same regime, mice proceeded to AA2 and showed the usual shift toward longer avoidance latencies, reflecting the emergence of caution when ITCs are punished (Fig. 6A, red). Again, no deficits were observed relative to controls.

**Figure 6.**
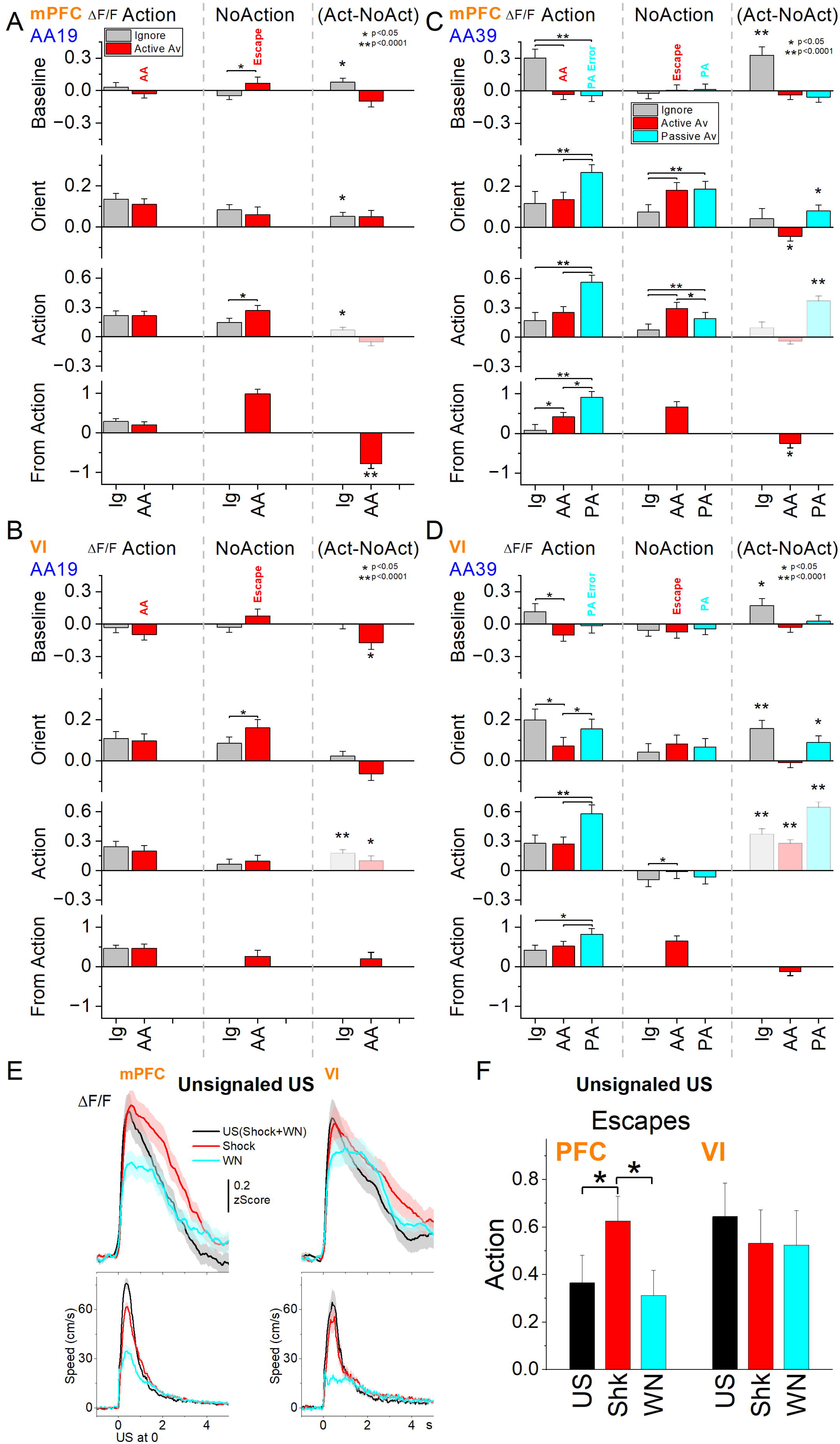
Optogenetic inhibition of mPFC neurons has little effect on signaled avoidance. ***A***, Effects of continuous green light at three different powers (10-30 mW) on learning AA1 (black circles) followed by AA2 (red circles) in naïve mice expressing eArch3.0 in mPFC neurons. In AA1, the CS signals the active avoidance contingency. In AA2, ITCs are punished. On every trial, light was delivered during the CS and US to inhibit mPFC activity. Mice learned and performed the task like control mice, which are not shown (but see D, E). During AA2, performance remained unchanged, although mice exhibited the typical shift toward longer avoidance latencies. Catch trials (NoCs) consisted of blank trials without light or consequence for crossing. ***B***, Same as A, but during subsequent AA3 task, in which CS1, CS1+Light, CS2, and CS2+Light trials were presented randomly. Light trials included the three powers used in A. CS1 signals the active avoidance (AA) contingency, whereas CS2 signals the passive avoidance (PA) contingency. ***C***, Optogenetic fiber placements in the mPFC and effect of blue light on neural activity in Vgat-Cre ChR2 mice. *Top left,* the schematic shows the range of depth placements tested in the mPFC of Vgat-Cre ChR2 mice in the sagittal plane (eArch3.0 fibers were around the dorsal portion of the range). *Bottom*, shows two examples of dorsal and ventral fibers from DAPI-stained sections. Example PSTH shows the effect of blue light in Vgat-Cre mice expressing ChR2 in GABAergic neurons recorded in freely behaving mice with an optrode implanted in mPFC. Continuous blue light for 1 s produces virtually complete sustained abolishment of multi-unit activity (MUA). The histogram is the average of 15 light trials. The same effects were observed in every mouse tested (n=3). ***D***, Effects of continuous blue light in mPFC on learning AA1 followed by AA2 in naïve mice expressing ChR2 in GABAergic neurons. The columns represent groups of mice with optical cannulas in mPFC or VI, with a third No Opsin group with optical fibers in mPFC that did not express ChR2. The AA1 and AA2 tasks are like those in A, including catch trials. The right panel shows the contrasts between AA1 and AA2 for each group. ***E***, Effect of blue light in the same three groups during AA3.

We next trained mice in AA3 tested mPFC inhibition, which includes four trial types (CS1, CS1+Light, CS2, CS2+Light) signaling active or passive responding. As in AA1 and AA2, inhibiting mPFC during CS1+Light trials had little effect on the probability of active avoidance across light powers (Fig. 6B). In contrast, passive avoidance was slightly but significantly impaired in CS2+Light trials at 10 mW (q(18)=5.25, p=0.0078) and 20 mW (q(18)=5.67, p=0.0041), with a trend at 30 mW (q(18)=3.96, p=0.052). However, the magnitude of impairment was modest (<15%). Thus, mPFC inhibition produced minimal effects on cued active responding and only a small decrement in passive responding.

The second optogenetic inhibition approach involved activating GABAergic neurons in Vgat-Cre ChR2 mice (cross of Vgat-Cre with Ai32 mice) to generate strong and sustained cortical inhibition, confirmed by optrode recordings (Fig. 6C, *top right*). In these mice, we applied blue light in mPFC (n = 13), or VI (n = 6). We also used No-Opsin controls (n = 10) that received the same light application but did not express ChR2. The large group size in mPFC is due to testing of a range of dorso-ventral fiber placements, to test for a regional difference (Fig. 6C top left and bottom). However, there were no differences between the mice based on fiber depth and the group was combined. In AA1, all groups learned at the same rate when every trial was a CS+Light trial (3 mW). Performance did not differ between the three groups (χ²(2) = 0.0094, p = 0.99). Notably, both mPFC and VI groups showed faster avoidance latencies than No-Opsin controls (mPFC: t(29) = 6.96, p < 0.0001; VI: t(31) = 4.66 p = 0.00011), an effect common across the cortical regions.

In AA2, all groups showed the expected increase in avoidance latencies as caution developed when ITCs were punished. In addition, No-Opsin controls showed the typical mild performance deficit (<10%) of transitioning to AA2, but this effect was absent in mPFC and VI groups, suggesting that cortical inhibition may act as a salient cue that facilitates responding, consistent with the earlier AA1 latencies.

In AA3, mice performed the four trial types after training. Light delivery during CS1 (active) trials produced little effect in any group and the latency shortening observed in AA1/2 for mPFC and VI groups was no longer apparent, suggesting that it is related to the initial learning phase. However, passive responding (CS2) was significantly, albeit very weakly (<15 %), impaired in both mPFC and VI groups (p < 0.0001), consistent with the effect also observed with eArch. No impairment occurred in the No-Opsin controls indicating that it is the cortical inhibition driving the mild impairment.

Together, the experiments show that mPFC is not required for generating cued actions under threat, even when mice must discriminate between cues and choose between action execution and deferment. Effects of mPFC inhibition were small, and largely shared by VI, indicating that the mPFC network is not necessary for learning or performing these goal-directed behaviors.

## Discussion

Our findings reveal how mPFC GABAergic neurons contribute to action selection under threat, clarifying long-standing questions about their role in cue processing, cautious responding, and outcome evaluation. Across a hierarchy of cued tasks, we found that much of the apparent task-related activity in both mPFC and VI could be explained by movement or baseline fluctuations, emphasizing the importance of controlling for these confounds when interpreting cortical signals. Yet after accounting for these factors, mPFC neurons—but not VI neurons—showed selective increases in activity during aversive outcomes, a signature of punished errors that may reflect the appraisal or salience of unexpected pain. Notably, simple avoidance contingencies were not encoded in mPFC population activity; instead, mPFC encoding emerged only under more demanding conditions in which mice had to discriminate cues and select between action and inaction. This pattern suggests that mPFC GABAergic neurons become differentially engaged when behavioral demands increase and cognitive load is high, rather than when the course of action is straightforward.

Single-cell imaging further revealed distinct GABAergic neuron subpopulations, some time-locked to cue onset, and others to the action itself. Classifying neurons based on their sensitivity to movement, revealed different classes of neurons encoding avoidance primarily during the challenging situation. This encoding was broad for passive avoidance but mostly restricted to a single class for active avoidance. Despite the rich activation related to cues, action selection, and punishment under threat, broad optogenetic inhibition of mPFC or VI minimally altered learning or behavioral performance, indicating that these signals do not appear necessary for executing cued actions under threat. Together, these results suggest that while mPFC neurons may evaluate and contextualize threat-related actions, they are not required to generate the actions themselves highlighting the distinction between encoding and controlling behavior.

### A valuable procedure for studying goal-directed contingencies

The sequence of tasks we employed provides a useful framework for probing how brain regions represent behavioral contingencies. In the initial noUS phase, mice generated spurious actions driven by baseline movement and cue salience. Once the active avoidance contingency was introduced in AA19, baseline movement and ITCs declined, and mice acquired selective, robust actions to the US-predictive CS1, with faster and more vigorous movement than spurious actions to non-predictive cues. Because CS1 can be systematically contrasted with CS2 and CS3, which remain non-predictive across these phases, and with itself in the noUS phase, the task structure provides controlled comparisons across contingencies and contexts. Introducing the passive avoidance contingency in AA39 with CS2 reduced overall action rates, including ITCs and correct actions to CS1, thereby increasing error rates relative to AA19. Action latencies also rose sharply, indicating hesitation and cautious responding as mice faced more demanding cue discrimination and action selection under threat. This design allowed CS2 responses in AA39 to be evaluated relative to their earlier non-predictive role in AA19 and contrasted with CS1, which continued to predict the active avoidance contingency, and CS3, which remained non-predictive throughout. Together, these behavioral findings show that mice rapidly learned distinct contingencies across progressively complex tasks and flexibly adapted their behavior as demands increased.

The combination of active and passive contingencies with multiple cues and intensities created a rich, systematic framework for dissecting neural computations underlying cue processing, decision-making, cautious action selection, and movement-related influences on neural activity.

### Cortical GABAergic neurons are movement-sensitive

Across all analyses, GABAergic neurons in mPFC and VI were strongly modulated by ongoing movement. Cross-correlation analyses showed that both translational and rotational components of movement correlated significantly with ΔF/F, with translational speed contributing most strongly. Directional analyses further suggested that mPFC GABAergic neurons do not encode head turn direction, whereas VI neurons show modest contraversive preference during ongoing movement.

In agreement with these results, every mixed-effects model used throughout this study—including those examining baseline, cue, action, and outcome trial phases—identified strong effects of movement covariates on cortical GABAergic neurons across all task conditions. Both baseline speed and window-specific speed were among the largest predictors of ΔF/F, often dwarfing the effects of the task variables themselves. This demonstrates that much of the apparent “task encoding” observed in raw traces reflects underlying movement coupling.

Together, these results indicate that cortical GABAergic neurons are intrinsically sensitive to movement, which covaries with behavioral states. As highlighted by many other studies (Musall et al., 2019; Steinmetz et al., 2019; Stringer et al., 2019; Zagha et al., 2022; Hormigo et al., 2023a), movement-related signals are sufficiently large and pervasive that careful statistical control is essential when evaluating whether neural activity genuinely reflects cue-, action-, or outcome-related processes.

### mPFC neurons encode punished errors independent of contingencies

A main finding across behavioral phases is that GABAergic neurons in mPFC consistently increase their activity during punishment in error trials. Although VI neurons also showed US-related activation, these signals were weaker than in mPFC. Thus, mPFC neurons displayed a selective sensitivity to aversive outcomes after controlling for baseline neural activity and movement. Several results converge on this interpretation. First, mPFC neurons showed systematic increases in ΔF/F during punished errors, including escapes in AA19 and passive-avoid errors in AA39. Second, mPFC neurons responded more strongly to footshock delivered alone than to the footshock embedded with auditory white noise (full US), suggesting that unexpected and concurrently unsignaled aversive events evoke stronger neural responses than aversive events accompanied by salient signals in another modality. This pattern aligns with human studies proposing that prefrontal circuits encode the appraisal or salience of pain (Lorenz et al., 2003; Lee et al., 2024). These effects were largely absent in VI, highlighting a regional dissociation in how aversive outcomes are represented. These findings indicate that mPFC GABAergic neurons exhibit selective, movement-independent encoding of punished errors, consistent with a role in evaluating aversive outcomes.

### mPFC encodes cued contingencies mostly in difficult situations

A second key finding is that mPFC encoding of avoidance contingencies emerges only under challenging conditions, not during simple, low-error tasks. During the easy AA19 task, mPFC neurons did not differentiate cues that predicted threat from those that did not, and activity during avoidance actions versus spurious actions showed little contingency-specific modulation beyond punishment-related activation. Also, the transition from noUS to AA19, where mice learn signaled active avoidance, did not reveal a selective change for the threat predictive cue, CS1, as similar changes occurred for the non-predictive cues.

In contrast, when the task became more demanding in AA39, mPFC activity was different for appropriate responses, such as active and passive avoids compared to spurious actions and ignored cues, respectively. This contingency-dependent recruitment manifested in several ways. mPFC neurons showed increased activity during active avoids only around the moment of action, and not during the CS, suggesting that mPFC becomes engaged around the time of action. This may reflect the appearance of cautious actions in the demanding AA39 environment, which are delayed but more vigorous. However, mPFC neurons also showed elevated activity during correct passive avoidance relative to correctly ignored trials, indicating recruitment during action withholding under threat. Importantly, since both action generation and deferment produced increased mPFC responses in AA39, this modulation may reflect increased cognitive load rather than contingency-related encoding. Thus, population-level GABAergic mPFC activity becomes engaged primarily under challenging conditions, marked by cue discrimination and hesitant action selection, consistent with a role in supporting higher cognitive load rather than directing the action itself.

### Selective encoding by populations of mPFC GABAergic neurons

Many studies examining mPFC activity under threat, across a wide range of behavioral contexts, report neural responses to cues, actions, and outcomes (e.g. (Simon et al., 2015; Diehl et al., 2018; Maggi et al., 2018; Jercog et al., 2021; Agetsuma et al., 2023; Ho et al., 2023; Martin-Fernandez et al., 2023; Sylte et al., 2024; Gu and Johansen, 2025; Lai et al., 2025)). In our study, we directly compared neural activity for equivalent cues that differed only in contingency (e.g., threat-predictive vs. non-predictive tones varying only in frequency), and for equivalent motor responses driven either by the predictive cue or by non-predictive cues (e.g., active avoids vs. spurious crossings; passive avoids vs. correctly ignored cues). We believe these comparisons provide a more selective assessment of task-specific encoding than other common contrasts with baseline activity or general movements (e.g., ITCs lack a cue altogether). Even under these stricter conditions, a subset of neurons showed selective activation linked to correct responding, consistent with the general view that some mPFC neurons encode action selection under threat.

When we classified individual mPFC GABAergic neurons using ΔF/F–movement cross-correlations, we found during AA19 that only a small subset exhibited selective encoding of correct actions. One subset (Class 3), which was weakly inhibited by movement, distinguished correct actions from spurious ones when aligned to CS onset but not when aligned to action execution. A second subset (Class 5), which was strongly movement-modulated, encoded correct actions only when aligned to the action itself, not the cue. The remaining classes were broadly responsive to the aversive outcome, underscoring the strong representation of foot-shock across the population. Thus, selective cue- and action-related encoding during simple contingencies was sparse and consistent with the predominant population-level patterns seen in photometry.

During the more challenging AA39 task, when mice learned the passive avoidance contingency, encoding patterns reorganized. Class 5 neurons continued to encode correct action generation and additionally represented correct action deferment. Class 1 and Class 3 also represented the correct passive responding compared to correctly ignored cues. Encoding of aversive outcomes remained widespread across classes, paralleling the photometry results.

When we clustered the movement profile of correct avoidance actions, different modes or styles were revealed including cautious responding where the mPFC activation was stronger. Further classifying the neural activity for each avoidance mode revealed two types of activations related to the cue and related to the action execution. The latter became more prominent during cautious responding, characterized by delayed but vigorous actions typical of high task difficulty.

Across task contingencies and increasing behavioral demands, mPFC GABAergic neurons integrated contextual, sensory, and motor information. Their selective engagement increased under high cognitive load and uncertainty, while broad activity reflected evaluation of aversive outcomes and the processing of cautious responding under threat.

### Is mPFC required for cued actions under threat?

Although mPFC involvement is frequently cited in cued actions under threat, including what is broadly termed “*avoidance*,” the term itself is used inconsistently in neuroscience and often refers to procedures that measure distinct constructs. For example, some studies use conditioned place preference/aversion, which assesses how animals evaluate contexts associated with reward or aversion rather than whether they can generate or withhold actions to avoid harm. Other studies use operant lever-press avoidance tasks in rodents, which are not species-specific defensive reactions like running away and have been shown to be essentially unlearnable (Bolles, 1970). Still others investigate conflict between approach and avoidance using a platform task (Diehl et al., 2018). In this context, conflict refers to the competition between opposing motivational demands, such as crossing a dangerous, busy road to reach a water fountain while thirsty, which is distinct from non-conflictual avoidance or approach (Miller, 1944; Botvinick et al., 2004; Aupperle et al., 2015). Finally, considering the species is also important because although mice and rats share threat-related behaviors they also exhibit differences (Blanchard et al., 2001, 2003, 2005).

Focusing specifically on classical active avoidance in rodents (Mowrer, 1939), lesion findings have been mixed. In rats, pre-training mPFC lesions slow learning whereas post-training lesions have no effect (Castro-Alamancos and Borrell, 1992). Another rat study reported similar slowing only when lesions included both dorsomedial and infralimbic mPFC (not dorsomedial alone), while post-training lesions were not examined (Moscarello and LeDoux, 2013). In mice, some studies report that dorsomedial mPFC inactivation impairs performance (Jercog et al., 2021), whereas others find minimal effects of irreversible lesions across avoidance contingencies (Zhou et al., 2022) in agreement with the optogenetic inactivation in the present study. In conflict procedures, there have also been inconsistencies even within the same study, such as muscimol inactivation delaying avoidance, while optogenetic inhibition is ineffective (Diehl et al., 2018).

In our study, action generation and deferment were never in direct conflict. We found that when baseline and movement variables are controlled in statistical models, encoding of avoidance contingencies is limited. Consistent with this sparse encoding, optogenetic inactivation of mPFC had minimal effects on avoidance in a variety of tasks of increasing difficulty.

We employed two different approaches to inhibit mPFC activity, including Vgat-ChR2 mice to activate GABAergic neurons, a well-established method for suppressing cortical activity via strong inhibition of local glutamatergic neurons. This manipulation produced robust suppression of cortical activity, and it did not impair avoidance behavior. These results are also consistent with previous studies showing that mPFC lesions do not impair avoidance behavior. Together, these findings indicate that avoidance behavior can be expressed in the absence of mPFC activity, consistent with a model in which other (cortical and/or subcortical) circuits normally mediate avoidance behaviors, while mPFC may contribute to other aspects of behavior.

## Materials and Methods

### Experimental Design and Statistical Analysis

The methods used in the present paper were like those employed in our previous studies (e.g., (Hormigo et al., 2023b; Zhou et al., 2024, 2026)). All procedures were reviewed and approved by the institutional animal care and use committee and conducted in adult (>8 weeks) male and female mice. Most experiments used a repeated-measures design in which each mouse or cell served as its own control (within-subject/cell comparisons), but we also compared experimental groups between subjects or cells (between-group comparisons). To test the main effects of experimental variables, we used either repeated-measures ANOVA or linear mixed-effects models.

Mixed-effects models were used to quantify relationships between ΔF/F activity (dependent variable) and experimental factors while accounting for repeated measures. Models included fixed effects for task contingency, outcome, cell/avoid classes, etc., and their interactions, along with window-specific covariates for head speed and baseline ΔF/F, both z-scored (mean = 0, SD = 1) within each analysis window. Random effects accounted for repeated measures by including sessions nested within subjects, or cells nested within subjects, depending on the analysis. Fixed effects included task-related factors and their interactions with speed, following the general form: ΔF/F ∼ (Factor1 * Factor2 *…) * Speed + (1 | Subject/Session) in lme4 (R package). Separate models were fit for each of the four behavioral windows, using mean measures for the baseline, orienting, avoidance, and from-action windows (normalized by window duration), with the latter windows corrected by the baseline. Generally, the baseline window (−0.5 to 0 s from CS onset) captures the pre-cue state at trial initiation, whereas the orienting window (0 to 0.5 s post-CS) reflects the initial response to the cue. The avoidance window (with task-specific durations) corresponds to the period when mice must generate or withhold a response to avoid punishment, excluding both the preceding orienting window and the subsequent escape interval. The from-action window aligns time series from the occurrence of a response (e.g., avoid, escape, passive avoid error), enabling comparison between avoidance responses occurring within the avoidance interval and escape responses in the later interval.

Including the window-specific head speed covariates, along with baseline ΔF/F and head speed measures, in the post-CS window models allowed us to dissociate movement-related modulation from neural encoding of task-related factors. Model fit was assessed by likelihood ratio tests comparing nested models, confirming that inclusion of covariates and interactions improved explanatory power. Model explanatory power was quantified using marginal and conditional R^2^ values (Nakagawa), corresponding to variance explained by fixed effects alone (R^2^_m_) and by the full model including random effects (R^2^_c_), computed from variance components of the fitted models (reported for each model). For significant main effects or interactions, post-hoc pairwise comparisons were performed using *emmeans* (R package) with Holm correction. For continuous covariates, slope effects were evaluated using *emtrends* (R package), with significance determined by t-tests (p < 0.05). Equivalence of movement measures was assessed using two one-sided tests (TOST) with a tolerance (delta) of ±6 cm/s (<50 % of baseline speed and <10% of peak speed) to confirm that classified neuronal activity differences were not generally attributable to movement differences.

To enable rigorous approaches, we maintain a centralized metadata system that logs all details about the experiments and is engaged for data analyses (Castro-Alamancos, 2022). Moreover, during daily behavioral sessions, computers run experiments automatically using preset parameters logged for reference during analysis. Analyses are performed using scripts that automate all aspects of data analysis from access to logged metadata and data files to population statistics and graph generation.

### Strains and Adeno-Associated Viruses (AAVs)

To record from GABAergic mPFC or VI neurons using calcium imaging, we injected a Cre-dependent AAV (AAV5-syn-FLEX-jGCaMP7f-WPRE (Addgene: 7×10^12^ vg/ml) in the mPFC or VI of Vgat-cre mice (Jax 028862; B6J.129S6(FVB)-Slc32a1^tm2(cre)Lowl^/MwarJ) to express GCaMP7f.

To excite GABAergic neurons in mPFC or VI using optogenetics, we crossed Vgat-cre and Ai32 (Jax 024109; B6.Cg-Gt(ROSA)26Sor^tm32(CAG-COP4*H134R/EYFP)Hze^/J) mice (Vgat-Chr2 mice). To broadly inhibit mPFC neurons using optogenetics, we expressed eArch3.0 by injecting AAV5-hSyn-eArch3.0-EYFP (UNC Vector Core, titers: 5.9×10^12^ vg/ml) in the mPFC of C57BL/6 mice (mPFC-Arch mice). The optogenetic methods used in the present study have been validated in previous studies using slice and/or in vivo electrophysiology (Hormigo et al., 2016, 2019, 2021). No-Opsin controls were injected with AAV8-hSyn-EGFP (Addgene, titers: 4.3×10^12^ GC/ml by quantitative PCR) or nil in the mPFC.

### Surgeries

Optogenetics and fiber photometry surgeries were performed under isoflurane anesthesia (∼1%), with postoperative carprofen administered for analgesia. AAVs (0.4 µl per site) were injected into the mPFC using stereotaxic coordinates given in millimeters relative to bregma (anteroposterior [AP], mediolateral [ML], dorsoventral [DV] below the bregma-lambda plane). For both Arch and GCaMP experiments, injections were made at AP = 2.0, ML = ±0.3, and DV = 1.5-2. Injections were unilateral for GCaMP and bilateral for Arch. Optical fibers were implanted at the time of surgery in the location of the AAV injections or centered on the midline: for fiber photometry, a 400 µm diameter fiber was placed unilaterally at ML = 0.3 mm; for miniscope recordings, a 600 µm diameter GRIN lens was placed unilaterally at ML = 0.3 mm; and for optogenetic experiments, a dual 200 µm diameter fiber was employed.

For optogenetics or fiber photometry experiments targeting VI, dual optogenetic cannulas or single fiber photometry implants were placed at AP = –2.6, ML = ±2.0, and DV = 0.2, with GCaMP AAV injections at the same coordinate as the cannula. Control animals without opsins (“No Opsin” mice) were implanted with cannulas in the mPFC or adjacent areas, and their data were pooled after confirming that light delivery produced similar behavioral effects across animals.

### Active Avoidance tasks

Mice are placed in a standard shuttle box (16.1″ x 6.5″) that has two compartments separated by a partition with side walls forming a doorway that the animal must traverse to shuttle between compartments. A speaker is placed on one side, but the sound fills the whole box and there is no difference in behavioral performance (signal detection and response) between sides. AA19 and AA39 sessions (n=7 each) are updated versions of previously described tasks (AA1 and AA3; e.g., (Zhou et al., 2022, 2023)) using three cues (CS1, CS2, CS3; 8, 4, 12 kHz tones) applied at three intensities (65-87 dB) to manipulate saliency and increase task difficulty. Thus, AA1 and AA3 are the same tasks as AA19 and AA39 using one or two salient cues respectively, while AA2 is like AA1 but punishes ITCs (0.2 s US).

During noUS sessions, the three CS were presented randomly without consequence during 7 sec windows, but crossing compartments (action) turned off the CS. During AA19, mice were presented with the same cues, but CS1 incorporated the active avoidance contingency. During AA39, the sessions continue as during AA19, but CS2 includes the passive avoidance contingency. In AA19/39, mice are free to cross between compartments during the intertrial interval; there is no consequence for intertrial crossings (ITCs).

CS1 trials in AA19/39 consist of a 7 sec avoidance interval followed by a 10 sec escape interval. During the avoidance interval, an auditory CS (8 kHz) is presented for the duration of the interval or until the animal produces a conditioned response (avoidance response) by moving to the adjacent compartment, whichever occurs first. If the animal avoids, by moving to the next compartment, the CS ends, the escape interval is not presented, and the trial terminates.

However, if the animal does not avoid, the escape interval ensues by presenting white noise and a mild scrambled electric foot-shock (0.3 mA) delivered through the grid floor of the occupied half of the shuttle box. This unconditioned stimulus (US) readily drives the animal to move to the adjacent compartment (escape response), at which point the US terminates, and the escape interval and the trial ends. Thus, an *avoidance response* will eliminate the imminent presentation of a harmful stimulus. An *escape response* is driven by presentation of the harmful stimulus to eliminate the harm it causes. Successful avoidance warrants the absence of harm. Each trial is followed by an intertrial interval (duration is randomly distributed; 25-45 sec range), during which the animal awaits the next trial.

In AA39, mice continue to be randomly presented the three cues but are subjected to a CS discrimination procedure in which they must respond differently to a CS1 and a CS2. Mice perform the basic signaled active avoidance to CS1 but also perform signaled passive avoidance to CS2. In AA39, if mice shuttle during the CS2 avoidance interval (7 sec), they receive a 0.5 sec foot-shock (0.3 mA) with white noise and the trial ends. If animals do not shuttle during the CS2 avoidance interval, the CS2 trial terminates at the end of the avoidance interval (i.e., successful signaled passive avoidance). AA1/2 and AA3 are the same tasks as AA19 and 39 using salient cues.

There are three main variables representing task performance. The percentage of active avoidance responses (% avoids) represents the trials in which the animal actively avoided the US in response to the CS. The response latency (latency) represents the time (sec) at which the animal enters the safe compartment after the CS onset; avoidance latency is the response latency only for successful active avoidance trials (excluding escape trials). The number of crossings during the intertrial interval (ITCs) represents random shuttling due to locomotor activity in the AA19 and AA39 procedures. The sound pressure level (SPL) of the auditory CS’s were measured using a microphone (PCB Piezotronics 377C01) and amplifier (x100) connected to a custom LabVIEW application that samples the stimulus within the shuttle cage as the microphone rotates driven by an actuator controlled by the application.

### Fiber photometry

We employed a 2-channel excitation (465 and 405 nm) and 2-channel emission (525 and 430 nm for GCaMP7f and other emissions) fiber photometry system (Doric Lenses). Alternating light pulses were delivered at 100 Hz (per each 10 ms, 465 is on for 3 ms, and 2 ms later 405 is on for 3 ms). While monitoring the 525 nm emission channel, we set the 465 light pulses in the 20-60 µW power range and then the power of the 405 light pulses was adjusted (20-50 µW) to approximately match the response evoked by the 465 pulses. During recordings, the emission peak signals evoked by the 465 (GCaMP7f) and 405 (isobestic) light pulses were acquired at 5-20 kHz and measured at the end of each pulse. To calculate F_o_, the measured peak emissions evoked by the 405 nm pulses were scaled to those evoked by the 465 pulses (F) using the slope of the linear fit. Finally, ΔF/F_o_ was calculated with the following formula: (F-F_o_)/F_o_ and converted to Z-scores. Due to the nature of the behavior studied, a swivel is essential. We employed a rotatory-assisted photometry system that has no light path interruptions (Doric Lenses). In addition, charcoal powder was mixed in the dental cement to ensure that ambient light was not leaking into the implant and reaching the optical fiber; this was tested in each animal by comparing fluorescence signals in the dark versus normal cage illumination.

### Miniscope imaging

We employed GRIN lenses (0.6mm diameter, 7.3mm length) and an nVista (Inscopix) recording system coupled with an electrical swivel. We added custom tungsten rods to the GRIN lenses to maximize the stability of the recordings. During each session, we adjusted the focus plane to record the best neurons, which were assigned as neurons per session. We used Inscopix Data Processing Software (IDPS) to extract ROIs from the raw miniscope movies. Briefly, the movies were preprocessed with a spatial bandpass filter, which removes the low and high spatial frequency content from the movies minimizing out of plane neuropil fluorescence and allows visual identification of putative neurons. The movies were then motion corrected, using the initial frame as the global reference. Finally, neurons were manually outlined on the processed movie, and the ΔF/F calcium activity was calculated from these ROIs. To assure stability during each recording session, the ROIs were visually verified from start to end for each video.

K-means clustering (scikit-learn) was performed on features extracted from ΔF/F (z-score) or movement (speed) time series traces. These features included peak amplitudes, mean amplitudes, and times to peak around specific events (e.g., turns, CS onset, or avoidance occurrence). Alternatively, clustering was also applied to principal component scores derived from the same traces, which were treated as an analog of spectral data for PCA.

### Optogenetics

The implanted optical fibers were connected to patch cables using sleeves. A black aluminum cap covered the head implant and completely blocked any light exiting at the ferrule’s junction. Furthermore, the experiments occurred in a brightly lit cage that made it difficult to detect any light escaping the implant. The other end of the patch cable was connected to a dual light swivel (Doric lenses) that was coupled to a green laser (520 nm; 100 mW) to activate Arch or a blue laser (450 nm; 80 mW) to activate ChR2. In experiments expressing Arch, Green light was applied continuously at different powers (10, 20, and 30 mW) during the indicated intervals. In experiments with ChR2 expression, blue light was delivered continuously at constant power (2-3 mW). Power is regularly measured by flashing the connecting patch cords onto a light sensor—with the sleeve on the ferrule.

During optogenetic experiments that involve avoidance procedures, we compared different trial types: CS and CS+Light. *CS trials* were standard avoidance trials specific to each procedure, without optogenetic stimulation. *CS+Light trials* were identical to CS trials, except that optogenetic light was delivered simultaneously with the CS and the US during the avoid and escape intervals. Catch trials present a blank no-CS trial without consequence to gauge random responding, which is also evaluated with ITCs. To perform within group repeated measures (RM) comparisons, the different trial types for a procedure were delivered randomly within the same session. In addition, the trials were compared between different groups, including No Opsin mice that did not express opsins but were subjected to the same trials including light delivery.

### Video tracking

All mice in the study (open field or shuttle box) were continuously video tracked (30-100 FPS) in synchrony with the procedures and other measures. During open field experiments, mice are placed in a circular open field (10″ diameter) that was illuminated from the bottom or in the standard shuttle box (16.1″ x 6.5″). We automatically tracked head movements with two color markers attached to the head connector –one located over the nose and the other between the ears. The coordinates from these markers form a line (head midline) that serves to derive several instantaneous movement measures per frame (Zhou et al., 2023). Overall head movement was separated into *rotational* and *translational* components (unless otherwise indicated, overall head movement is presented for simplicity and brevity, but the different components were analyzed). Rotational movement was the angle formed by the head midline between succeeding video frames multiplied by the radius. Translational movement resulted from the sum of linear (forward vs backward) and sideways movements. *Linear* movement was the distance moved by the ears marker between succeeding frames multiplied by the cosine of the angle formed by the line between these succeeding ear points and the head midline. *Sideways* movement was calculated as linear movement, but the sine was used instead of the cosine. Pixel measures were converted to metric units using calibration and expressed as speed (cm/sec). We used the time series to extract window measurements around events (e.g., CS presentations). Measurements were obtained from single trial traces and/or from traces averaged over a session. In addition, we obtained the direction of the rotational movement with a *Head Angle* or *bias* measure, which was the accumulated change in angle of the head per frame (versus the previous frame) zeroed by the frame preceding the stimulus onset or event (this is equivalent to the rotational speed movement in degrees). For all signals, including neural activity, peaks correspond to the signed *extrema* (maximum or minimum) within each analysis window. Window responses were also quantified as the mean of the time series within each window. The *time to peak* is when the *extrema* occurs versus event onset.

To detect spontaneous turns or movements from the head tracking, we applied a local maximum algorithm to the continuous head angle or speed measure, respectively. Every point is checked to determine if it is the maximum or minimum among the points in a range of 0.5 sec before and after the point. A change in angle of this point >10 degrees was a detected turn in the direction of the sign. We further sorted detected turns or movements based on the timing of previous detected events.

### Histology

Mice were deeply anesthetized with an overdose of isoflurane. Upon losing all responsiveness to a strong tail pinch, the animals were decapitated, and the brains were rapidly extracted and placed in fixative. The brains were sectioned (100 µm sections) in the coronal or sagittal planes. All sections were mounted on slides, cover slipped with DAPI mounting media, and all the sections were imaged using a slide scanner (Leica Thunder). We used an APP we developed with OriginLab (Brain Atlas Analyzer) to align the sections with the Allen Brain Atlas Common Coordinate Framework (CCF) v3 (Wang et al., 2020). This reveals the location of probes and fluorophores versus the delimited atlas areas.

Figure legends are also inserted in the Results section to facilitate perusal.

## Acknowledgments

Supported by NIH grants to MAC.

---

**Figure 1-Supplement 1.**
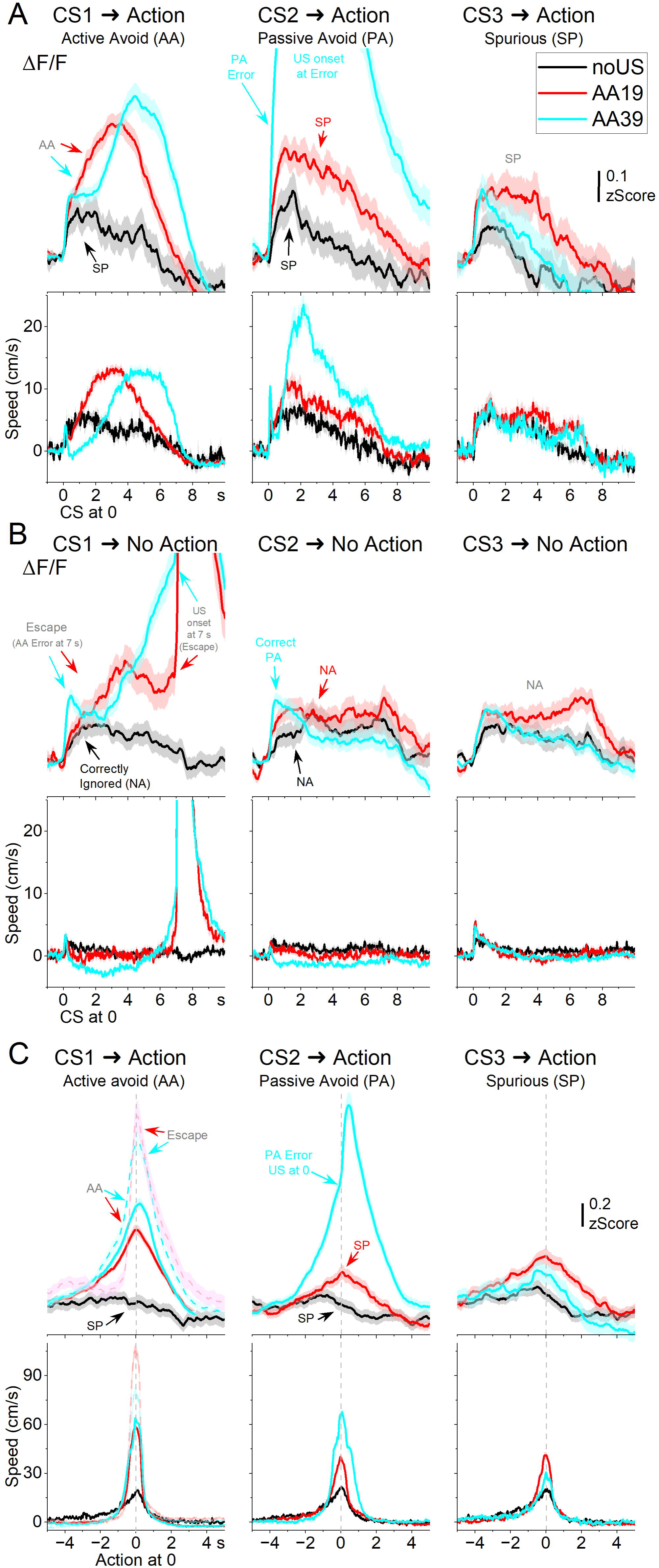
mPFC and VI GABAergic neurons differentially encode exploratory movement turning direction. ***A***, ΔF/F calcium imaging, overall movement, rotational movement, and angle of turning direction for detected movements classified by the turning direction (ipsiversive and contraversive; red and cyan; red is dashed to improve visualization of overlapping traces) versus the side of the recording (implanted optical fiber) in mPFC. Around zero time, the animals spontaneously turn their head in the indicated direction (zero time is the peak speed of the turn, which is used for alignment). The columns show all turns (left), those that included no turn peaks 3 s prior (middle), and peaks selected at intervals >5 s (right). Note that the speed of the movements was similar in both directions (the y-axis speed is truncated to show the rising phase of the movement). ***B***, Same as A for VI. ***C***, Population measures (mean, peak amplitude, and time to peak for traces 3 s around the detected peaks) of ΔF/F and overall movement for the different classified peaks in mPFC. Asterisks denote significant differences (p<0.05) between ipsiversive and contraversive movements. In the mPFC, neuronal activity did not differ between ipsiversive and contraversive turns across any condition (all turns: t(583)= 0.43 p=0.66; no turns 3 sec prior: t(583)= 0.35 p=0.73; turns per >5 sec: t(1157)= 1.5 p=0.13). ***D***, Same as C for VI. VI neuron activity showed modest direction selectivity. Contraversive movements elicited significantly greater ΔF/F responses for all turns (t(583)= 3.6 p=0.0004), isolated turns per >5 sec (t(1157)= 3.31 p=0.0009), but not for turns following quiescence (no turns 3 sec prior: t(583)= 1.8 p=0.07). (n=7 mice in mPFC, n=5 mice in VI)

**Figure 2-Supplement 1.**
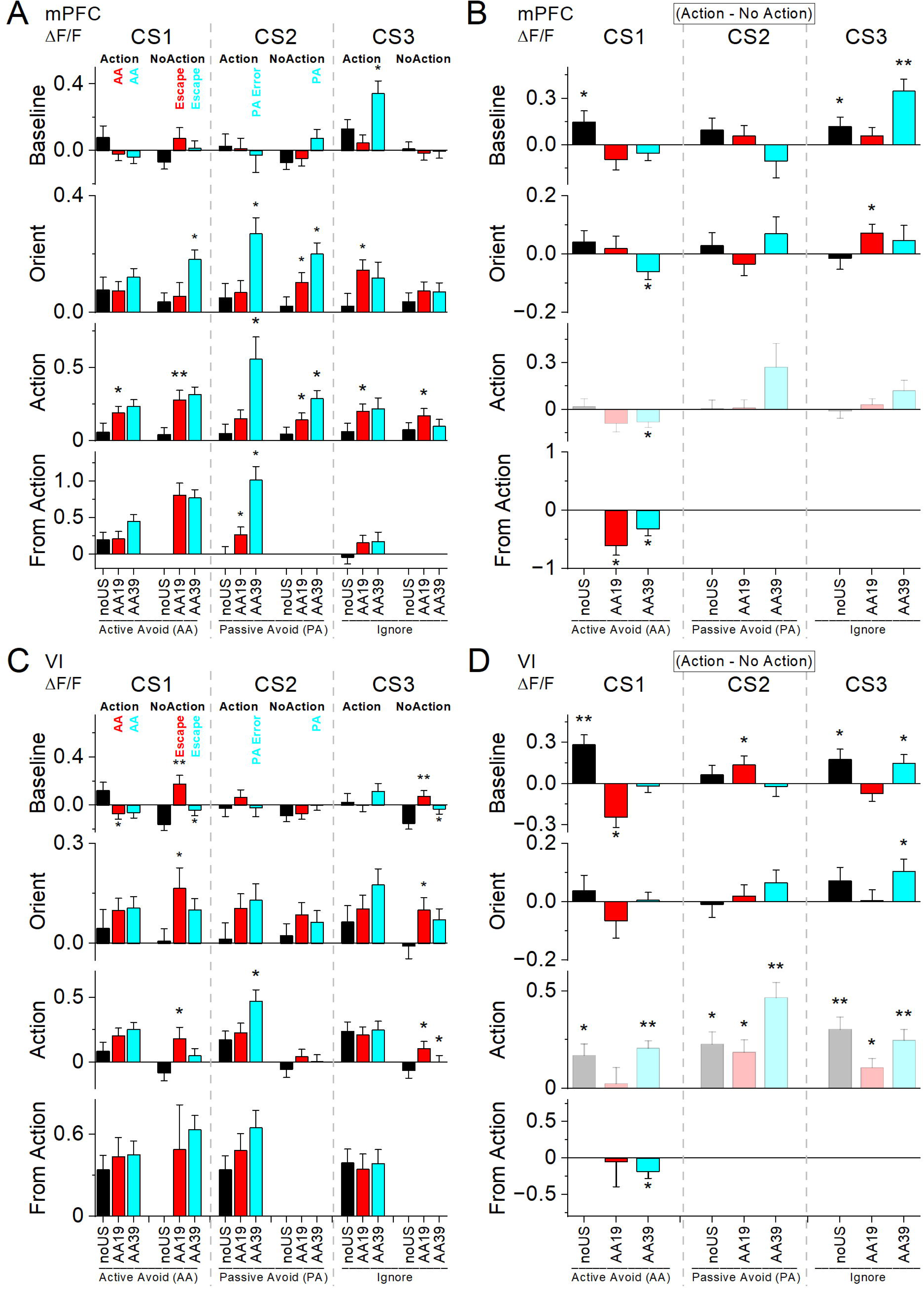
**Movement traces (overall speed)** across task phases (noUS, AA19, AA39), cues (CS1–CS3), and cue intensities (low, medium, high), separated by Action (***A***) and No-Action (***B***) outcomes for all mPFC and VI mice. Dashed boxes indicate correct or error responses for the active avoidance (AA) and passive avoidance (PA) contingencies; traces outside the boxes correspond to spurious actions or correctly ignored cues for the Ig contingency. Note that escapes are AA errors. Each panel overlays the three cue intensities. Some traces are intentionally truncated to highlight specific features, but the full trace is shown in C. ***C***, Movement for the Action trials in A, replotted aligned to the action occurrence (From-Action is defined as the time at which the animal crosses the compartments door). Values represent averages across mice.

**Figure 2-Supplement 2.**
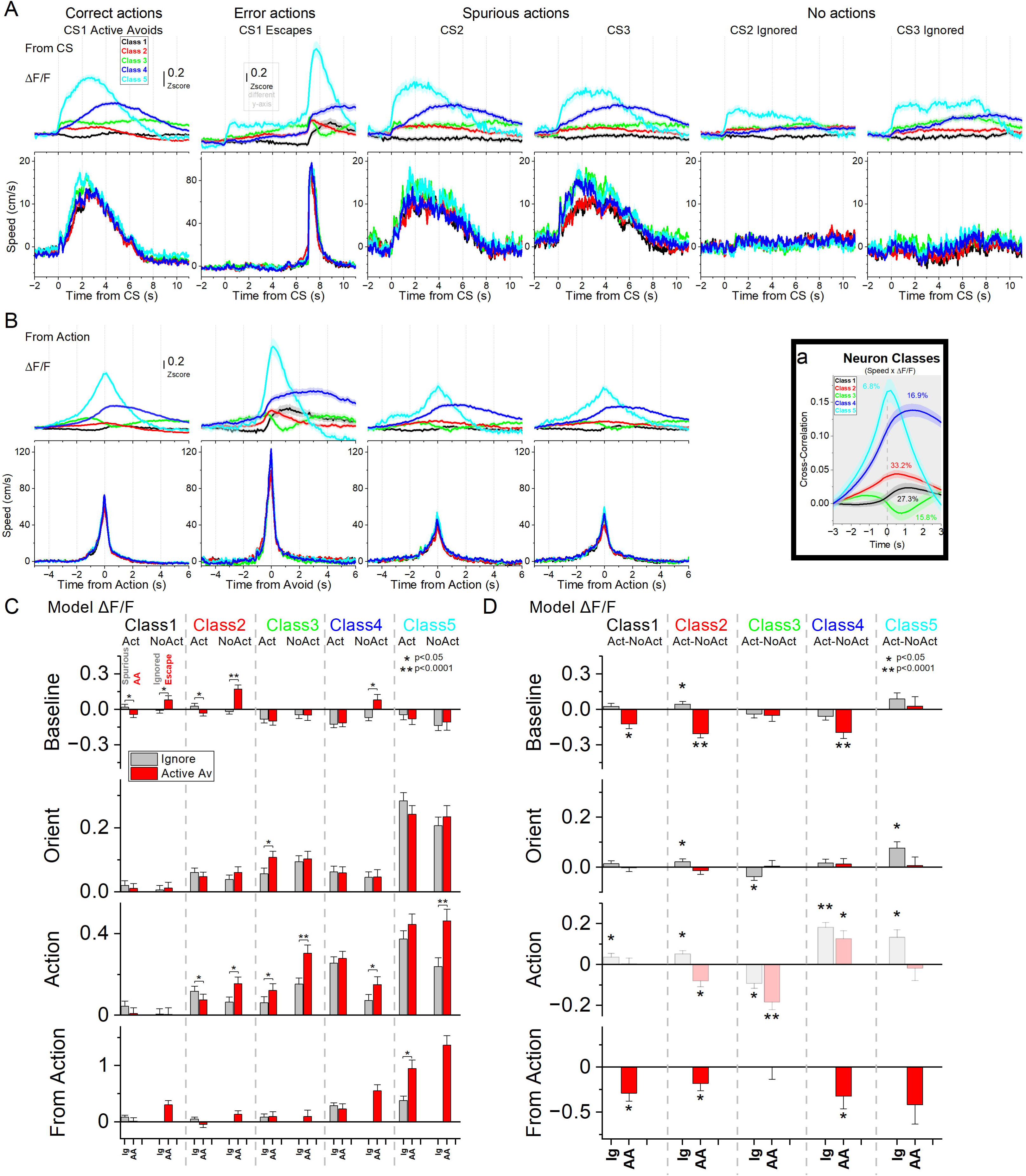
Marginal means of movement from mixed-effects models. Population-level marginal means of movement speed estimated from linear mixed-effects models across all mice. Left panels (***A***) compare trial types across task phases; right panels (***B***) compare Action versus No-Action outcomes within each task and CS. Asterisks in left panels denote significant differences relative to the preceding task phase. Asterisks in right panels denote significant differences between Action and No-Action outcomes. Each panel row corresponds to one model window: baseline (pre-CS), orienting (0–0.5 s post-CS), action (0.5–7 s post-CS), and from-action (−2 to 2 s relative to crossing). Movement speed is expressed in cm/s (averaged within each time window). **Baseline window**. Baseline movement differed across task phases and action outcomes. There were significant main effects of Task (χ²(2) = 52.0, p < 0.0001) and Outcome (χ²(1) = 199.9, p < 0.0001), as well as CS × Outcome (χ²(2) = 13.9, p = 0.001) and Task × Outcome (χ²(2) = 37.5, p < 0.0001) interactions. During noUS, baseline speed was higher on Action than No-Action trials across all CSs (t(3052) > 7, p < 0.0001). In AA19, baseline speed was higher for Action trials during CS2 (t(3082) = 4.44, p < 0.0001) and CS3 (t(3078) = 3.74, p = 0.00018), but not during CS1 (t(3116) = 0.38, p = 0.7). In AA39, baseline speed differed between Action and No-Action trials for CS3 (t(3096) = 7.43, p < 0.0001), CS2 (t(3098) = 4.1, p < 0.0001), and CS1 (t(3067) = 2.4, p = 0.015). Baseline movement decreased from noUS to AA19 (t(239) = 5.76, p < 0.0001) but did not differ between AA19 and AA39 (t(246) = 1.26, p = 0.2). (R^2^_m_ = 0.18; R^2^_c_ = 0.5) **Orienting window**. Orienting responses varied as a function of task and tone intensity. A significant Task × Tone Intensity interaction was observed (χ²(4) = 25.5, p < 0.0001), with a Task × Tone Intensity × CS interaction (χ²(8) = 16.2, p = 0.04). At high intensity, orienting responses increased from noUS to AA19 for CS1 (t(3071) = 3.21, p = 0.0039) and CS3 (t(2861) = 3.4, p = 0.0019), but not for CS2 or at lower intensities. (R^2^_m_ = 0.18; R^2^_c_ = 0.22) **Action window**. Movement during the action window showed main effects of CS (χ²(2) = 25.7, p < 0.0001) and tone intensity (χ²(2) = 73.4, p < 0.0001). Significant CS × Task (χ²(4) = 26.0, p < 0.0001) and CS × Tone Intensity (χ²(4) = 35.8, p < 0.0001) interactions were observed. Movement differed strongly between Action and No-Action trials (χ²(1) = 1274, p < 0.0001), with significant interactions between Outcome, CS, Task, and Tone Intensity (all p < 0.0001). (R^2^_m_ = 0.41; R^2^_c_ = 0.47) **From-Action window**. From-Action is defined as the time at which the animal crosses the door. Movement aligned to the action event varied by CS (χ²(2) = 135.84, p < 0.0001), Task (χ²(2) = 165.79, p < 0.0001), and Tone Intensity (χ²(2) = 39.89, p < 0.0001). A CS × Task interaction was present (χ²(4) = 259.42, p < 0.0001), along with a CS × Task × Tone Intensity interaction (χ²(8) = 18.05, p = 0.02). (R^2^_m_ = 0.72; R^2^_c_ = 0.77) **noUS Action trials**. During noUS, movement did not differ among CSs when intensities were pooled. At high intensity, CS2 differed from other CSs during noUS but not during AA19. **AA19 Action trials**. During AA19, movement during CS1 Action trials was greater than during noUS spurious actions (t(1871) = 8.82, p < 0.0001). Smaller increases were observed for CS2 (t(2284) = 4.2, p < 0.0001) and CS3 (t(2048) = 3.31, p = 0.0028), restricted to medium and high intensities. Within AA19, CS1 Action trials differed from CS2 (t(3102) = 3.14, p = 0.0033) and CS3 (t(3090) = 3.92, p = 0.00027). **AA39 Action trials**. In AA39, movement during CS2 Action trials differed from CS2 Action trials in AA19 (t(2653) = 5.91, p < 0.0001), with effects strongest at medium and high intensities. Movement during CS1 Action trials did not differ between AA19 and AA39, whereas CS3 Action trials decreased relative to AA19 (t(2381) = 3.22, p = 0.0028). Within AA39, CS2 Action trials differed from CS1 Action trials (t(3136) = 5.6, p < 0.0001). Comparable effects were observed in from-action–aligned analyses. **AA39 No-Action trials**. During AA39, movement during No-Action CS2 trials differed from No-Action CS2 trials in AA19 (t(1710) = 2.96, p = 0.0062). A similar difference was observed for CS1 No-Action trials (t(2408) = 2.63, p = 0.017), whereas no change was observed for CS3 No-Action trials (t(1466) = 0.23, p = 0.81).

**Figure 3-Supplement 1.**
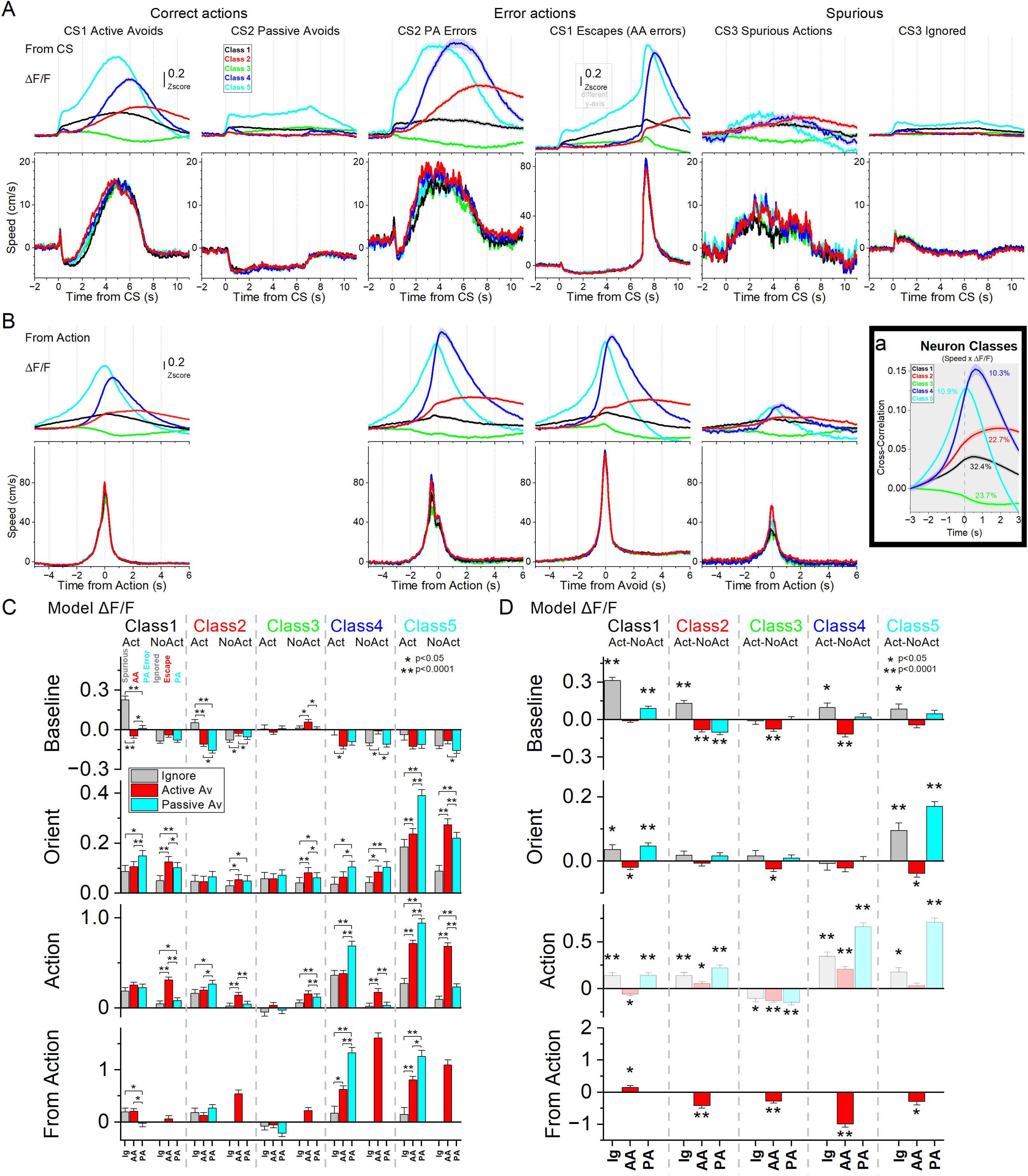
**ΔF/F fiber photometry traces and corresponding movement traces for mPFC mice** across task phases (noUS, AA19, AA39) and cues (CS1–CS3), separated by Action (***A***) and No-Action (***B***) outcomes. For each cue (CS1, CS2, CS3; left, middle, right panels), traces from the three task phases are overlaid. Movement traces for each condition are shown in the lower panels. Arrows indicate correct or error responses for the active avoidance (AA) and passive avoidance (PA) contingencies. Some traces are intentionally truncated to highlight specific features, but the full trace is shown in C. ***C***, Traces for the Action trials in A, replotted aligned to the action occurrence (From-Action is defined as the time at which the animal crosses the door).

**Figure 3-Supplement 2.**
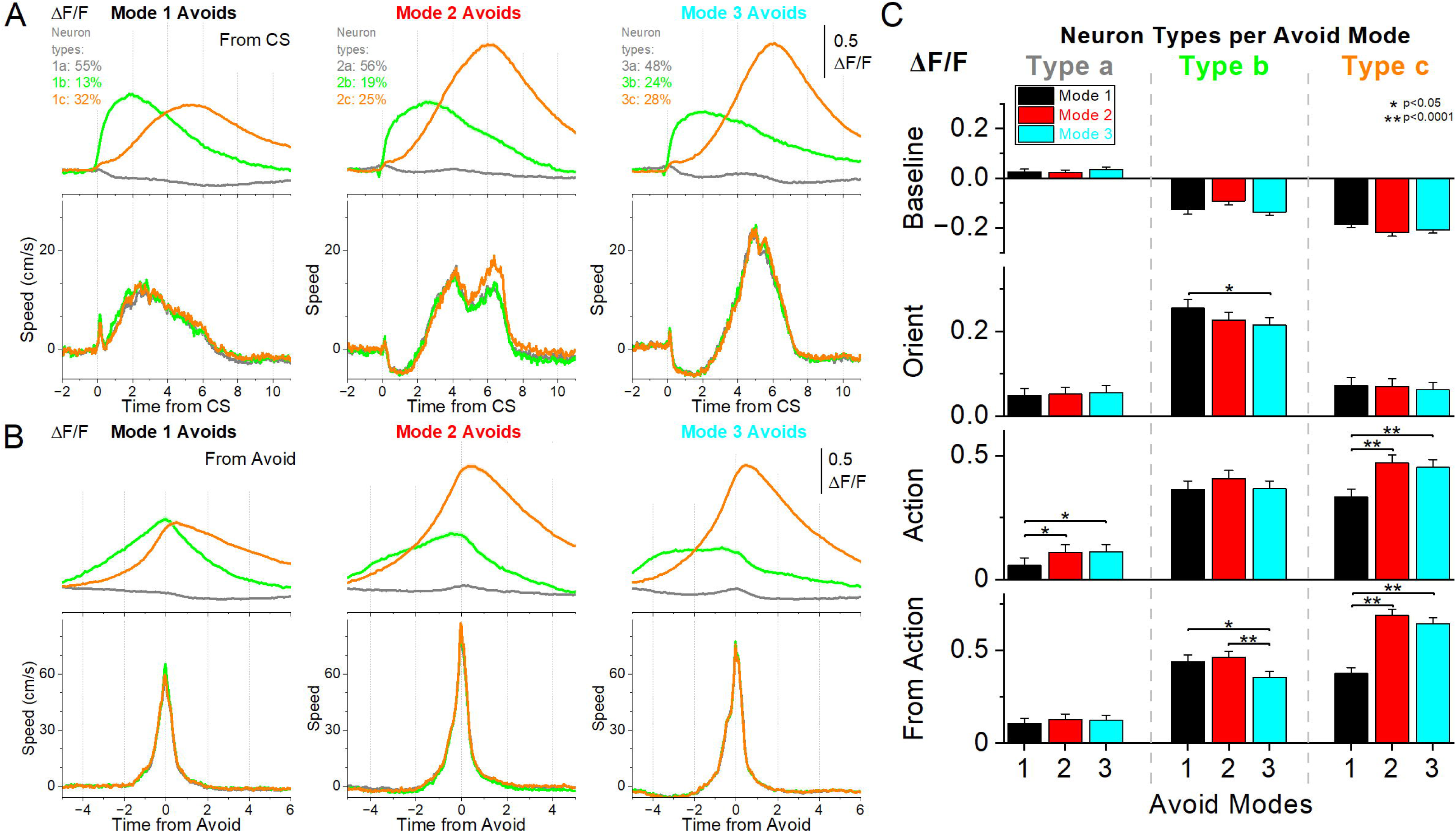
Full mixed-effects models of ΔF/F activity across task phases controlling for movement and baseline activity. Population estimates of marginal means from full mixed-effects models of ΔF/F fiber-photometry signals are shown for mPFC (***A****, **B***) and VI (***C****, **D***) GABAergic neurons. Models include all task phases (noUS, AA19, AA39) and incorporate covariates for window-specific movement, baseline movement, and baseline ΔF/F (excluded from the baseline window). Random effects were sessions nested within mice. Left panels (***A****, **C***) show comparisons across task phases for each CS and outcome; right panels (***B***, ***D***) show comparisons between Action and No-Action outcomes within each task and CS. Asterisks in left panels denote significant differences relative to the preceding task phase; asterisks in right panels denote significant differences between Action and No-Action outcomes. Rows correspond to the baseline, orienting, action (CS-aligned), and from-action analysis windows (From-Action is defined as the time at which the animal crosses the door). Shaded bars in the action window indicate that movement was controlled within, but not between, outcomes. Actions not classified as AA, PA, Escape, or PA Error are spurious actions. **Baseline window**. Baseline ΔF/F was strongly influenced by baseline movement (χ²(1) = 318.62, p < 0.0001). After controlling for movement, a significant Task × CS × Outcome interaction was observed (χ²(4) = 26.16, p < 0.0001), while the four-way Task × CS × Outcome × Group interaction was marginal (χ²(4) = 8.18, p = 0.085). Between-group contrasts revealed higher baseline ΔF/F in mPFC compared to VI for CS3 during noUS spurious-action trials (t(266) = 2.27, p = 0.024) and No-Action trials (t(50) = 2.48, p = 0.016). Within mPFC, baseline ΔF/F differed between Action and No-Action trials for CS1 and more prominently for CS3 during AA39. Across tasks, only CS3 showed a significant increase in baseline ΔF/F for Action trials from AA19 to AA39. In VI, baseline ΔF/F was higher for Action than No-Action trials across multiple CSs and task phases, including CS1 in noUS, CS2 in AA19, and CS3 in noUS and AA39. This relationship reversed for CS1 in AA19, where higher baseline VI activity preceded escapes (t(3086) = 3.36, p = 0.00078). Baseline ΔF/F changed in opposite directions for active avoids and escapes across AA19. (R^2^_m_ = 0.24; R^2^_c_ = 0.27) **Orienting window**. Orienting-related ΔF/F was significantly modulated by orienting movement (χ²(1) = 96.09, p < 0.0001), baseline movement (χ²(1) = 44.08, p < 0.0001), and baseline ΔF/F (χ²(1) = 401.6, p < 0.0001). After accounting for these covariates, the Task × CS × Outcome interaction was marginal (χ²(4) = 8.87, p = 0.061), and the corresponding four-way interaction including Group was not significant (χ²(4) = 5.05, p = 0.28). Between-group contrasts revealed higher orienting ΔF/F in mPFC than VI for CS2 during AA39 correct passive avoids (t(33) = 2.64, p = 0.012). In mPFC, orienting ΔF/F differed between escapes and active avoids for CS1 and between spurious-action and No-Action trials for CS3 in AA19. Across tasks, orienting ΔF/F increased for CS2 No-Action and CS3 spurious-action trials from noUS to AA19, and for CS1 escapes and CS2 Action and No-Action trials from AA19 to AA39. In VI, significant outcome-related differences were limited to CS3 during AA39, and task-related increases were observed only for CS1 and CS3 No-Action trials from noUS to AA19. (R^2^_m_ = 0.32; R^2^_c_ = 0.48) **Action window**. ΔF/F during the action window was strongly influenced by action-window movement (χ²(1) = 306.78, p < 0.0001), baseline movement (χ²(1) = 196.08, p < 0.0001), and baseline ΔF/F (χ²(1) = 1874.7, p < 0.0001). After controlling for movement within outcome categories, a significant Task × CS × Outcome interaction was observed (χ²(4) = 10.16, p = 0.037), along with a significant Task × CS × Outcome × Group interaction (χ²(4) = 10.88, p = 0.027). Between-group contrasts showed higher action-window ΔF/F in mPFC than VI for CS2 correct passive avoids (t(23) = 3.65, p = 0.0013) and CS1 escapes (t(22) = 3.50, p = 0.002) during AA39. In mPFC, ΔF/F increased across task phases for CS1 as active avoids and escapes emerged between noUS and AA19, and for CS2 with the emergence of passive avoid outcomes in AA39. CS3 also showed increases for both Action and No-Action trials from noUS to AA19. In VI, ΔF/F increases were observed for CS1 with the emergence of escapes in AA19 and for CS2 with the emergence of passive avoid errors in AA39, while CS3 showed increases primarily in No-Action trials. Differences between Action and No-Action trials were larger in VI than mPFC in this window. (R^2^_m_ = 0.58; R^2^_c_ = 0.69) **From-Action window.** From-action ΔF/F was significantly modulated by from-action movement (χ²(1) = 154.73, p < 0.0001), baseline movement (χ²(1) = 40.47, p < 0.0001), and baseline ΔF/F (χ²(1) = 96.45, p < 0.0001). After controlling for these covariates, significant Task × CS × Outcome (χ²(4) = 18.09, p = 0.0019) and Task × CS × Group interactions were observed (χ²(4) = 22.65, p = 0.00015), with a marginal four-way interaction (χ²(4) = 9.6, p = 0.047). Between-group contrasts revealed higher from-action ΔF/F in VI than mPFC for CS2 (t(47) = 2.35, p = 0.022) and CS3 (t(38) = 3.20, p = 0.0027) spurious actions during noUS. In mPFC, from-action ΔF/F was higher for escapes than active avoids in AA trials during both AA19 and AA39. ΔF/F also increased for CS2 spurious actions from noUS to AA19 and for CS2 passive avoid errors in AA39 relative to spurious actions in AA19. In VI, outcome-related differences in the from-action window were limited, with enhanced activation during escapes observed only in AA39. (R^2^_m_ = 0.51; R^2^_c_ = 0.66)

**Figure 4-Supplement 1.**
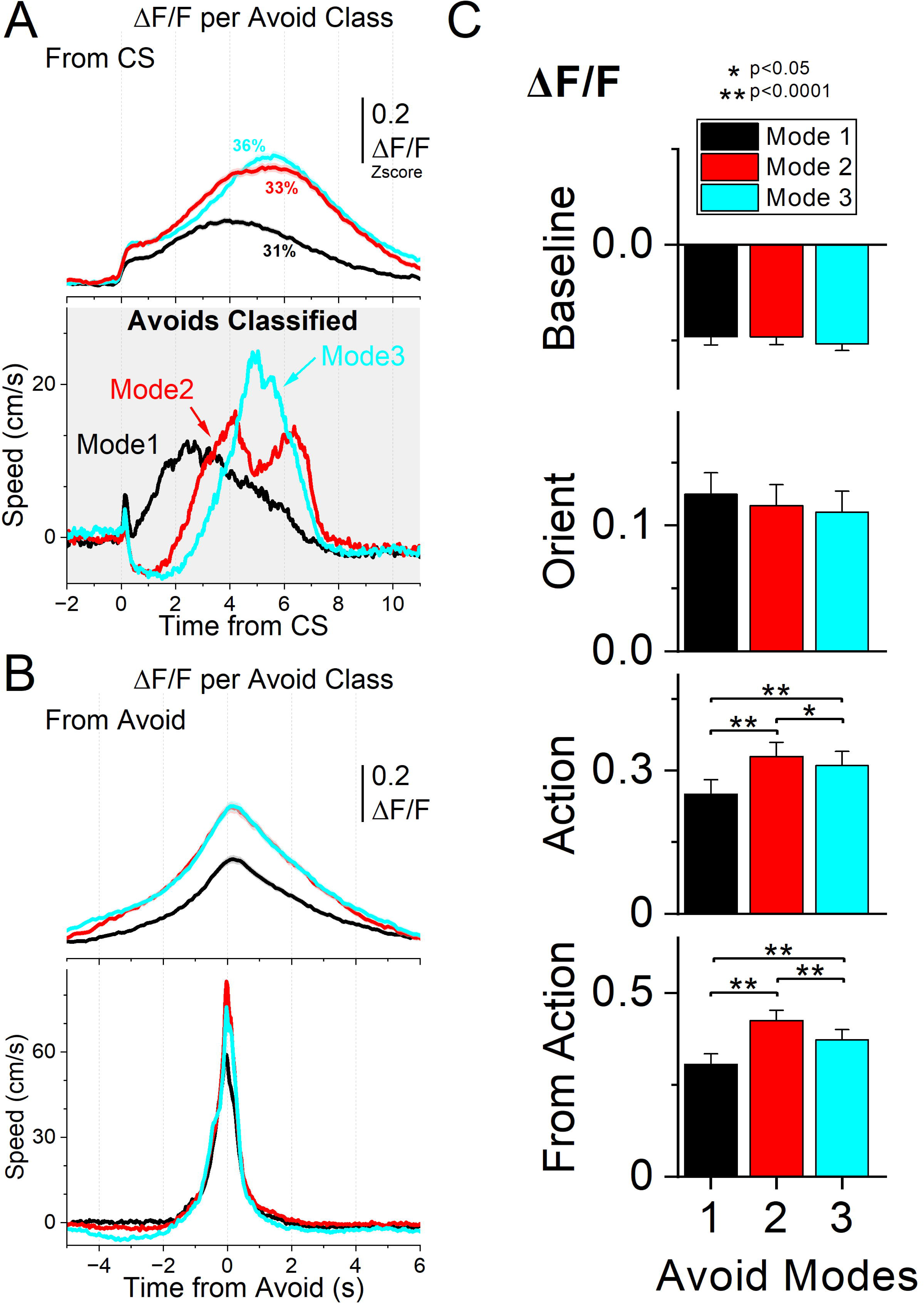
Movement-related neuron classes during a difficult avoidance context (AA39 task). ***A***, ΔF/F traces (top) and movement traces (bottom) for five mPFC GABAergic neuron classes identified by k-means clustering of cross-correlation functions between speed and ΔF/F (inset **a**). Class 1 (black) and Class 2 (red) neurons show moderate positive correlations with movement. Class 3 (green) neurons show negative correlations with movement. Class 4 (blue) and Class 5 (cyan) neurons show strong positive correlations, with Class 5 activity preceding Class 4 around zero lag. Traces are aligned to CS onset and shown separately for CS1–CS3 and behavioral outcome (action vs. No-Action), which may reflect correct responses, errors, or spurious actions depending on contingency. In AA39, CS1 signals active avoidance and an action is the correct response, whereas CS2 signals passive avoidance and an action is an error. ***B***, Same as A, aligned from-action (defined as the time at which the animal crosses the door) for Action outcomes. Panels for correct CS2 passive avoids are not shown because these trials do not include an action and therefore cannot be aligned to the from-action window. ***C***, Marginal mean ΔF/F estimates from linear mixed-effects models for the baseline, orienting, action, and from-action windows, comparing Ignore (Ig), Active Avoid (AA), and Passive Avoid (PA) contingencies across outcomes (action vs. No-Action) for each neuron class. ***D***, Same as C, but comparing outcomes within each contingency for each neuron class. Transparency in the action window indicates that movement was not controlled between outcomes in that model. **Baseline window**. The mixed-effects model revealed a significant Contingency × Outcome interaction (χ²(1) = 194.01, p < 0.0001) and a Contingency × Outcome × Class interaction (χ²(4) = 66.83, p < 0.0001). (R^2^_m_ = 0.08; R^2^_c_ = 0.17) **Orienting window**. A significant Contingency × Outcome interaction (χ²(2) = 106.17, p < 0.0001) and a significant Contingency × Outcome × Class interaction (χ²(8) = 81.89, p < 0.0001) were observed. (R^2^_m_ = 0.13; R^2^_c_ = 0.48) **Action window**. The model revealed a significant Contingency × Outcome interaction (χ²(2) = 179.6, p < 0.0001) and a significant Contingency × Outcome × Class interaction (χ²(8) = 240.7, p < 0.0001). (R^2^_m_ = 0.33; R^2^_c_ = 0.57) **From-action window**. A significant Contingency × Outcome × Class interaction was observed (χ²(8) = 257.09, p < 0.0001). (R^2^_m_ = 0.33; R^2^_c_ = 0.56)

**Figure 5-Supplement 1.**
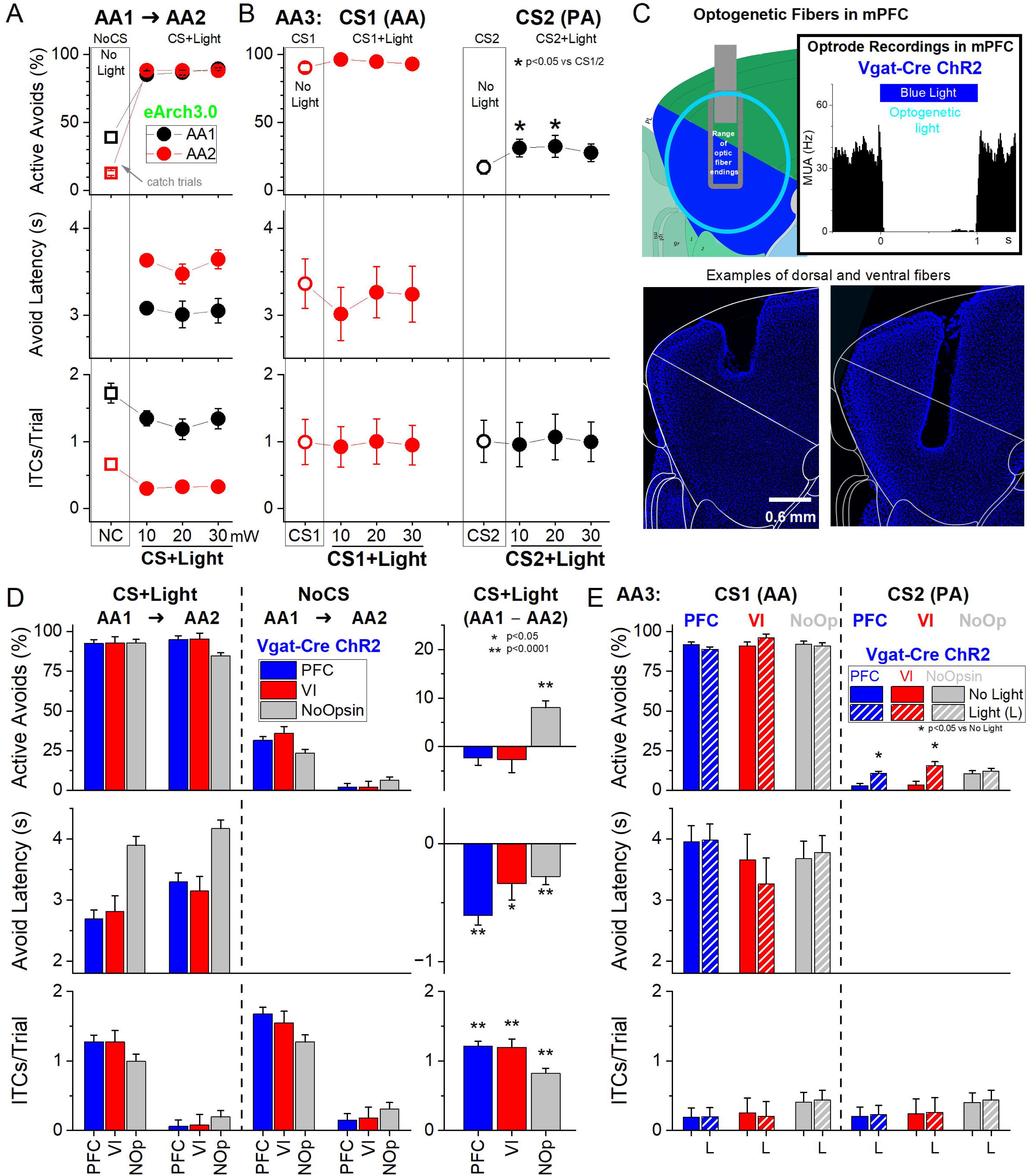
Activation of mPFC GABAergic neurons across three distinct signaled active avoidance modes. ***A***, k-means clustering of movement speed time-series from CS onset (gray panel) revealed three distinct active avoidance modes. Mode 1 avoids were initiated earlier after CS onset, whereas Mode 2 and Mode 3 avoids were delayed and faster (peak speed vs Mode 1: p < 0.0001), reflecting increasingly cautious responding. The top panel shows the average ΔF/F activity of all recorded mPFC neurons for each avoidance mode. mPFC activation was weaker during Mode 1, compared to Mode 2 and Mode 3. ***B***, Same as A, aligned from-action (avoid) occurrence. ***C***, Marginal means (ΔF/F) from the linear mixed-effect models for the baseline, orienting, avoidance, and from-action windows for the three active avoidance Modes. **Action window**. Mode 2 (t(4912) = 6.37, p < 0.0001) and Mode 3 (t(5246) = 4.86, p < 0.0001) avoids exhibited higher activation than Mode 1 avoids, and Mode 2 tended to have higher activation than Mode 3 avoids (t(4698) = 2.04, p = 0.04). **Fron-Action window**. Mode 2 vs Mode 1: t (4955) = 7.85, p < 0.0001; Mode 3 vs Mode 1: t(5236) = 4.27, p < 0.0001; Mode 2 vs Mode 3: t(4441) = 4.29, p < 0.0001.

